# Characterization of enhancer fragments in *Drosophila robo2*

**DOI:** 10.1101/2022.08.01.502399

**Authors:** Gina Hauptman, Marie C. Reichert, Muna A. Abdal Rhida, Timothy A. Evans

## Abstract

Receptor proteins of the Roundabout (Robo) family regulate axon guidance decisions during nervous system development. Among the three *Drosophila robo* family genes *(robo1, robo2,* and *robo3*), *robo2* displays a dynamic expression pattern and regulates multiple axon guidance outcomes, including preventing midline crossing in some axons, promoting midline crossing in others, forming lateral longitudinal axon pathways, and regulating motor axon guidance. The identity and location of enhancer elements regulating *robo2’s* complex and dynamic expression pattern in different neural cell types are unknown. Here, we characterize a set of 17 transgenic lines expressing GAL4 under the control of DNA sequences derived from noncoding regions in and around *robo2,* to identify enhancers controlling specific aspects of *robo2* expression in the embryonic ventral nerve cord. We identify individual fragments that confer expression in specific cell types where *robo2* is known to function, including early pioneer neurons, midline glia, and lateral longitudinal neurons. Our results indicate that *robo2*’s dynamic expression pattern is specified by a combination of enhancer elements that are active in different subsets of cells. We show that *robo2*’s expression in lateral longitudinal axons represents two genetically separable subsets of neurons, and compare their axon projections with each other and with Fasciclin II (FasII), a commonly used marker of longitudinal axon pathways. In addition, we provide a general description of each fragment’s expression in embryonic tissues outside of the nervous system, to serve as a resource for other researchers who may be interested in *robo2* expression and its functional roles outside the CNS.

## Introduction

The *Drosophila lea/robo2* gene encodes a transmembrane protein (Robo2) that is a member of the evolutionarily conserved Roundabout (Robo) family of axon guidance receptors. Robo receptors canonically regulate midline crossing of axons by signaling midline repulsion in response to Slit ligands. In addition to their evolutionarily conserved role in Slit-dependent midline repulsion, individual Robo family members also regulate other developmental processes both inside and outside of the nervous system. In *Drosophila,* three Robo family members are present (Robo1, Robo2, and Robo3) and each plays a distinct subset of developmental roles that depend on differences in expression pattern, differences in protein activity, or both (Rajagopalan *et al.* 2000a; b; Simpson *et al.* 2000a; b; Spitzweck *et al.* 2010; Evans and Bashaw 2010; Evans *et al.* 2015).

*Drosophila robo2* was initially named *leak (lea)* and was first identified by virtue of its head development defect (“head broad”) in Nüsslein-Volhard and Wieschaus’s foundational genetic screen for mutations affecting the patterning of the larval cuticle (Nüsslein-Volhard *et al.* 1984). Mutations in *lea* were later found to fail to complement mutations in *robo2* (Schimmelpfeng *et al.* 2001). *robo2* is expressed in a dynamic pattern during *Drosophila* embryonic ventral nerve cord development. Its expression is broad in early pioneer neurons, where it is expressed alongside *robo1* during the early stages of axon guidance, and becomes more restricted as the nerve cord develops (Rajagopalan *et al.* 2000a; Simpson *et al.* 2000a). In late embryonic development, Robo2 protein in the nerve cord is largely restricted to longitudinal axons in the lateral-most region of the neuropile (Rajagopalan *et al.* 2000b; Simpson *et al.* 2000b), and *robo2* mRNA remains detectable in motor neurons (in particular, ventrally-projecting RP motor neurons) through stage 17 (Santiago *et al.* 2014).

Outside of the developing ventral nerve cord, Robo2 embryonic expression (protein and/or mRNA) has been reported in the anterior ectoderm during head involution (Schimmelpfeng *et al.* 2001), thoracic visceral mesoderm and chordotonal neurons (Kraut and Zinn 2004), ectodermal stripes (Rajagopalan *et al.* 2000a; Ordan and Volk 2015), developing heart/pericardial cells (Simpson *et al.* 2000a; Santiago-Martínez *et al.* 2006), tracheal branches (Simpson *et al.* 2000a; Englund *et al.* 2002; Parsons *et al.* 2003), ventral longitudinal muscles (Simpson *et al.* 2000a; Kramer *et al.* 2001; Santiago *et al.* 2014), and the embryonic gonad (Weyers *et al.* 2011). Additional aspects of *robo2* expression have been reported in post-embryonic stages, including expression in the adult posterior midgut (Biteau and Jasper 2014), glial cells in the larval leg imaginal disc (Sasse and Klämbt 2016), adult testis (Stine *et al.* 2014), and motor neurons and gustatory neurons in the adult leg (Brierley *et al.* 2009; Mellert *et al.* 2010). While some links have been established between aspects of *robo2* expression and specific transcription factor proteins, for example Hb9’s role in regulating *robo2* transcription in RP motor neurons (Santiago *et al.* 2014), the location or sequence of individual enhancer elements in *robo2* have not been identified for any of the subsets of cells in which *robo2* is expressed.

Here, we examine the embryonic expression patterns of 17 *Drosophila* transgenic lines expressing the GAL4 transcriptional activator under the control of genomic DNA fragments derived from intergenic and intron sequences in and around the *robo2* gene (Pfeiffer *et al.* 2008; Jenett *et al.* 2012). We crossed each line to a *UAS-TauMycGFP* reporter strain and we have characterized the GFP expression pattern conferred by each fragment in the embryonic ventral nerve cord during the major stages of axon guidance (stages 12 through 17). We identify individual fragments that confer expression in specific cell types where *robo2* is known to function, including early pioneer neurons, midline glia, and lateral longitudinal neurons. Our results indicate that *robo2’s* dynamic expression during embryonic CNS development is specified by a combination of enhancer elements located both upstream of the *robo2* promoter and within the large *robo2* first intron that are active in different subsets of cells. We also show that *robo2’s* expression in lateral longitudinal axons represents at least two genetically separable subsets of commissural neurons with cell body positions and axon projection patterns that are morphologically distinct from each other. In addition, we provide a general description of expression characteristics for each fragment in embryonic tissues outside of the nervous system, to serve as a resource for other researchers who may be interested in *robo2* expression and its functional roles outside of the CNS.

## Results

To identify potential enhancer sequences regulating *robo2* transcription during embryonic ventral nerve cord development, we took advantage of a series of transgenic strains generated as part of a large scale effort to identify *Drosophila* enhancers expressed in neuronal subsets in the adult brain (FlyLight collection, https://flweb.janelia.org/cgi-bin/flew.cgi) (Jenett *et al.* 2012). These transgenic lines each carry a small segment of DNA (average size approx. 3,000 bp) derived from introns or upstream intergenic regions of genes known to be expressed in the brain, cloned upstream of the GAL4 transcriptional activator coding sequence (Pfeiffer *et al.* 2008). We obtained 17 of these GAL4 lines carrying sequences derived from in or around the *robo2* gene **(Figure 1; Table 1).** We crossed each of these lines to a reporter line carrying a GAL4-responsive *UAS-TauMycGFP (UAS-TMG)* transgene, and used an anti-GFP antibody to examine the expression of each GAL4 line throughout embryogenesis in the progeny of each cross. The TauMycGFP (TMG) reporter protein binds to microtubules and labels the cell bodies and axons of neurons in which GAL4 is expressed.

**Figure 1.**
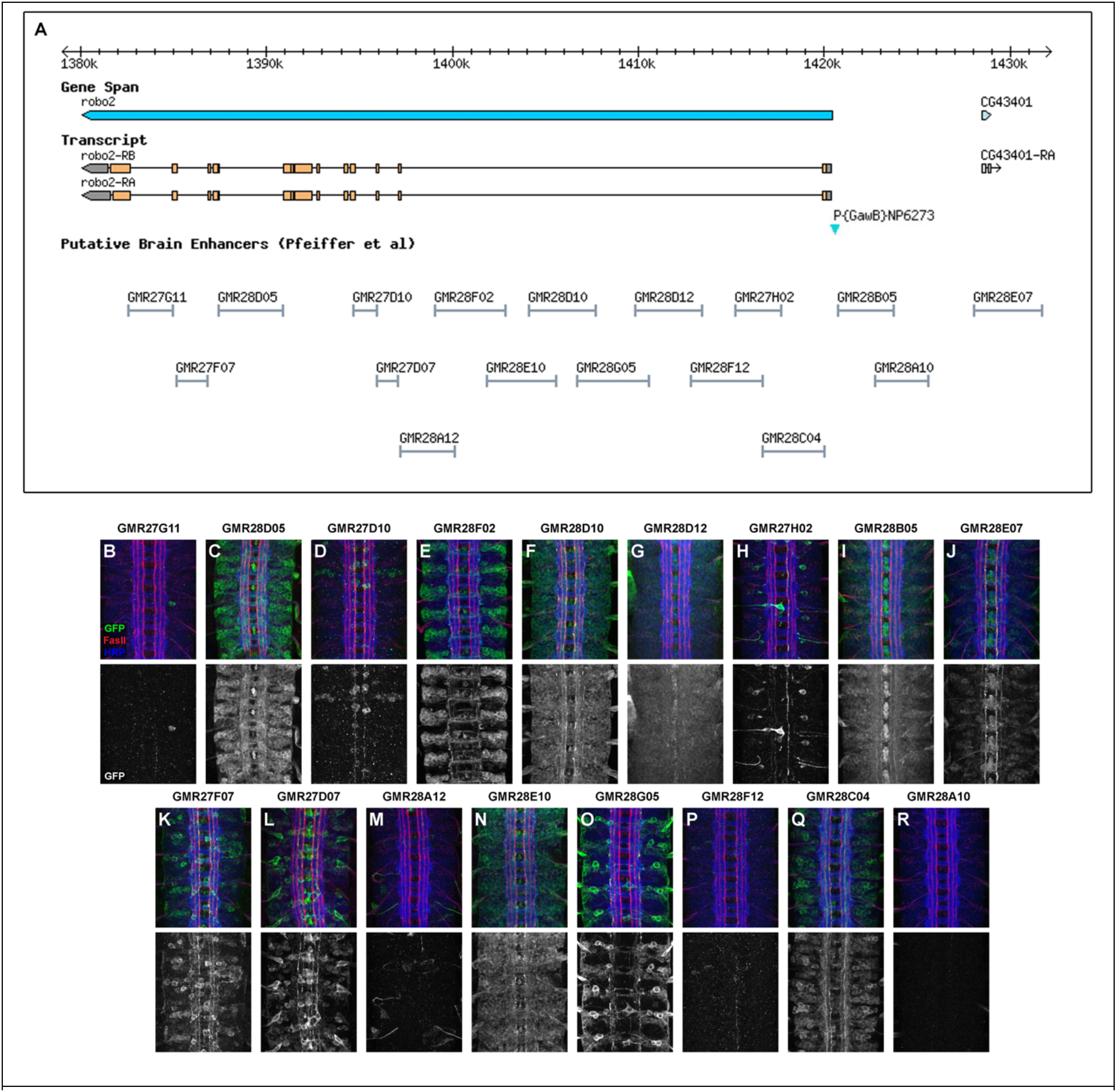
*robo2* enhancer fragments and summary of GAL4 transgene expression patterns. (A) Schematic of the *robo2/lea* genomic region illustrating the intron/exon structure of the *robo2* transcription unit (grey boxes, 5’ and 3’UTR; orange boxes, coding exons). Transcription of *robo2* proceeds from right to left. Two *robo2* transcripts with identical sequence are reported *(robo2-RA* and *robo2-RB).* Positions of putative enhancer fragments used to create GAL4 transgenic lines by Pfeiffer et al (Pfeiffer *et al.* 2008) are shown below (“Putative Brain Enhancers”). Screenshot taken from http://flybase.org/cgi-bin/gbrowse2/dmel/?Search=1;name=FBgn0002543. The location of the GAL4 enhancer trap transgene *P{GawB}NP6273 (robo2^GAL4^)* is indicated by the cyan triangle. The insertion site is 217 bp upstream of the predicted *robo2* transcriptional start site, and does not overlap with the GMR28C04 or GMR28B05 fragments. (B-R) GAL4 expression patterns in stage 16-17 ventral nerve cords from embryos carrying the indicated GAL4 transgenes along with *UAS-TauMycGFP (UAS-TMG),* labeled with antibodies against GFP (green; labels GAL4-expressing cells), FasII (red; labels a subset of longitudinal axon pathways), and HRP (blue; labels all axons). Lower panels show anti-GFP channel alone.

**Table 1.**
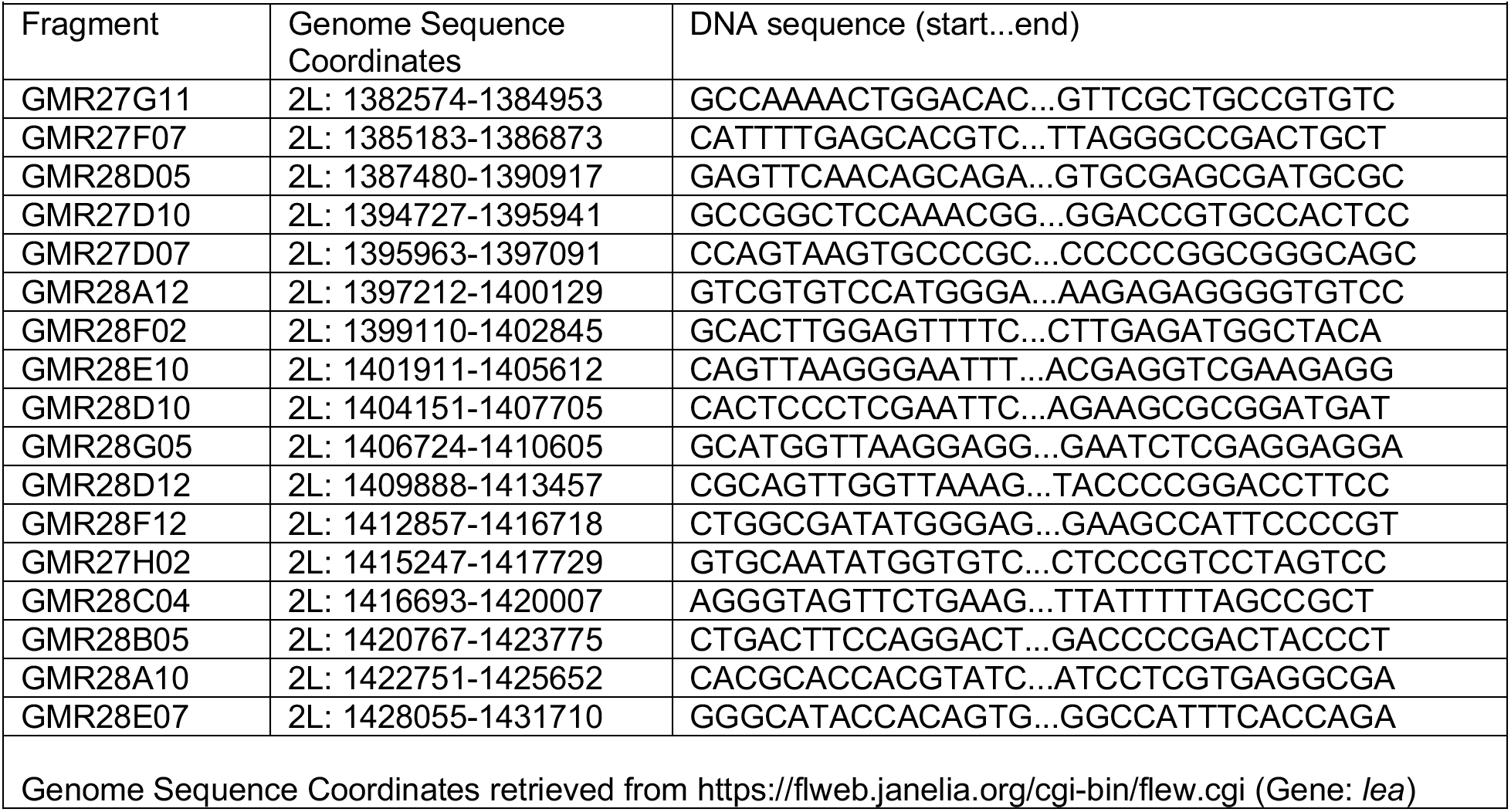
DNA sequence fragments from the *Drosophila* GAL4 lines used in this study.

We first examined the expression of each GAL4 line in whole-mount embryos stained with an anti-GFP primary antibody followed by a horseradish peroxidase (HRP)-conjugated secondary antibody, and developed with 3,3-Diaminobenzidine (DAB), which produces a brown precipitate in the presence of HRP. We examined and photographed whole embryos under a dissection microscope and noted general aspects of GFP expression throughout the embryo (e.g. brain, VNC, gut, epidermis, muscle, salivary glands) from the initial stages of axon outgrowth through the end of embryogenesis (embryonic stages 12-17).

Then, to more closely examine the expression of each GAL4 line in the ventral nerve cord during embryonic development, we performed immunofluorescence staining with an anti-GFP primary antibody in combination with antibodies against HRP (anti-HRP), which crossreacts with a pan-neural epitope in *Drosophila* and labels all of the axons in the embryonic ventral nerve cord (Jan and Jan 1982; Haase *et al.* 2001), and Fasciclin II (anti-FasII), which labels pioneer axons during the early stages of axon guidance and a subset of mature longitudinal axon pathways in older embryos (Grenningloh *et al.* 1991). We dissected nerve cords from these fluorescently-stained embryos and collected confocal z-stack images of nerve cords at each developmental stage from 12-17.

In the following sections, we first provide an overview of the embryonic expression patterns conferred by each of the 17 putative enhancer fragments, compared with endogenous Robo2 expression and a *robo2^GAL4^* enhancer trap line, followed by a more in-depth description of fragments of particular interest which direct expression in neuronal and glial subsets in which Robo2 is known to be functionally important, including lateral longitudinal neurons and midline glia.

### Endogenous Robo2 protein expression in the embryo

Robo2 protein is broadly expressed in CNS neurons beginning at stage 12, when early extending axons pioneer the initial longitudinal and commissural axon pathways (Rajagopalan *et al.* 2000a; b; Simpson *et al.* 2000a). *robo2* mRNA is detectable slightly earlier, at stage 11 (Simpson *et al.* 2000b). Robo2 protein becomes more restricted as the ventral nerve cord develops, and at later stages is detectable only on longitudinal axons in the most lateral region of the neuropile (Rajagopalan *et al.* 2000b; Simpson *et al.* 2000b). We and others have shown that anti-HA staining of modified *robo2* loci including an N-terminal 4xHA tag reproduces the pattern seen with antibodies against Robo2 protein in the late-stage embryonic CNS (Spitzweck *et al.* 2010; Howard *et al.* 2021). In whole *robo2^HA-robo2^* embryos stained with anti-HA, Robo2 protein is also clearly detectable in the gut, head, and epidermis in addition to its CNS expression **(Figure 2A).** We used a similar engineered *robo2* allele with an N-terminal 4xMyc tag to detect Robo2 protein expression in embryos co-labeled with anti-HRP (which labels all axons in the embryonic CNS) and anti-FasII (which labels motor axons and a subset of longitudinal axon pathways in the ventral nerve cord) from stages 12-17, which illustrates Robo2’s early broad expression and later restriction to lateral axons **(Figure 2B-G).** Like Robo1, Robo2 protein is cleared from commissural segments of midline-crossing contralateral axons (Rajagopalan *et al.* 2000a), so it is not clear from examining endogenous Robo2 protein which Robo2-expressing axons are ipsilateral and which (if any) are contralateral.

**Figure 2.**
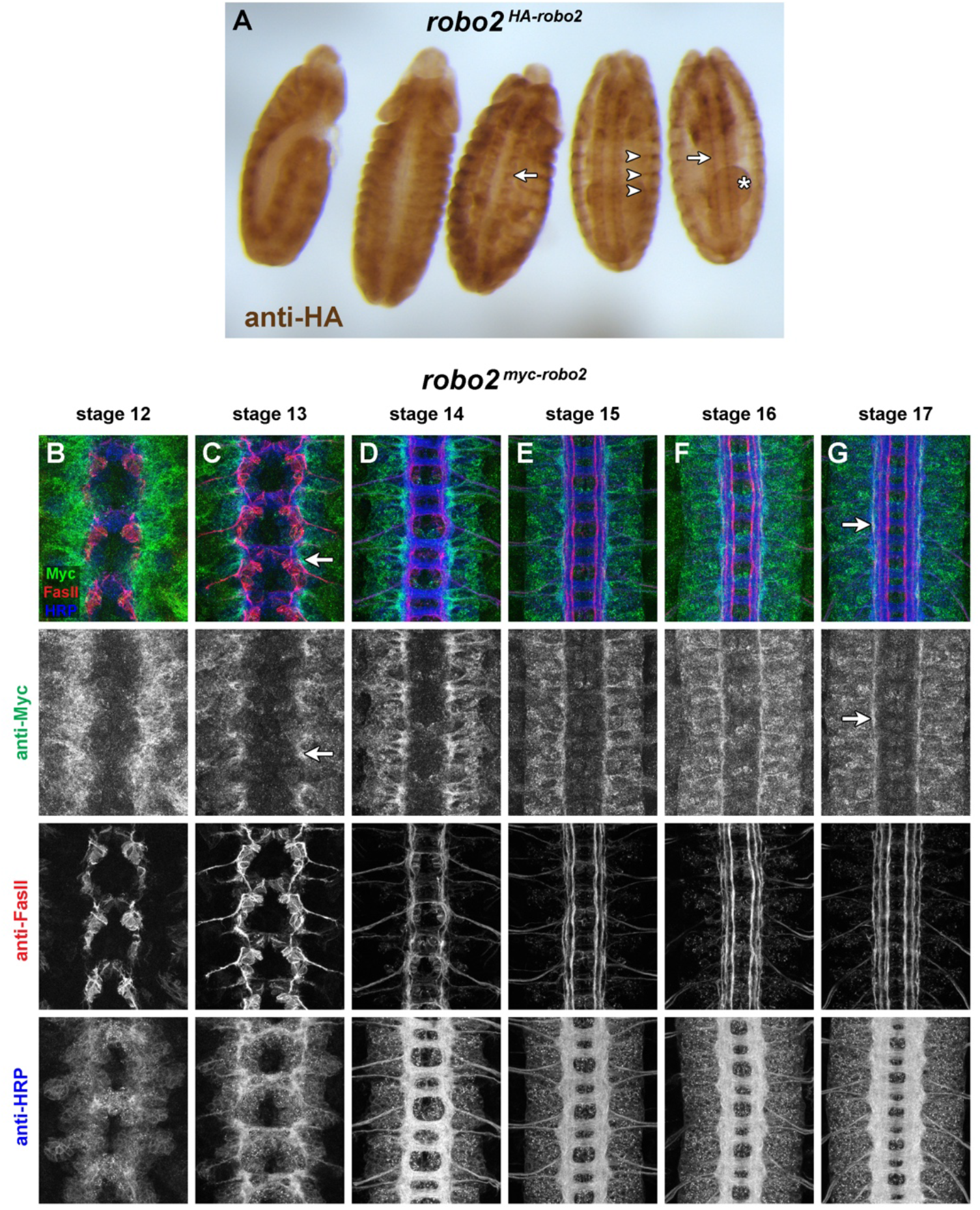
Endogenous expression of Robo2 protein. (A) Embryos homozygous for an HA-tagged *robo2^HA-robo2^* allele stained with an anti-HA antibody (brown). Representative embryos from various developmental stages are arranged in order of increasing age from left to right. Lateral view of youngest embryo on left; all others ventral side up. Robo2 protein is detectable throughout the ventral nerve cord in early stages and on lateral longitudinal axons at late stages (arrows). Outside of the CNS, Robo2 protein is also readily detectable in ectodermal stripes (arrowheads) and in the gut (asterisk). (B-G) Ventral nerve cords from stage 12-17 embryos homozygous for a myc-tagged *robo2^myc-robo2^* allele stained with anti-myc (green), anti-Fasll (red) and anti-HRP (blue) antibodies. Lower images show isolated channels for each of the three antibodies. Robo2 protein is broadly expressed in neurons during the early stages of axon outgrowth and detectable on axons and growth cones of pioneer neurons (C, arrow), and by late stages is mainly detectable on longitudinal axons within the lateral-most one-third of the neuropile (G, arrow).

### Embryonic expression pattern of a *robo2^GAL4^* enhancer trap insertion

*P{GawB}NP6273* is a GAL4 enhancer trap transgene insertion located 217 bp upstream of the predicted *robo2* transcriptional start site, recovered in a large-scale screen for novel insertions of the GAL4 enhancer trap element P{GawB} (Hayashi *et al.* 2002). GAL4 expression in this line recapitulates aspects of the *robo2* expression pattern including midline glia, early pioneer neurons and lateral longitudinal neurons (Evans *et al.* 2015). In whole embryos carrying *robo2^GAL4^* and *UAS-TMG,* GFP is strongly and broadly expressed throughout the embryonic epidermis. Although this makes it difficult to detect internal aspects of GFP expression in these intact embryos, some internal GFP can clearly be seen, including strong midline expression in the ventral nerve cord **(Figure 3A).**

**Figure 3.**
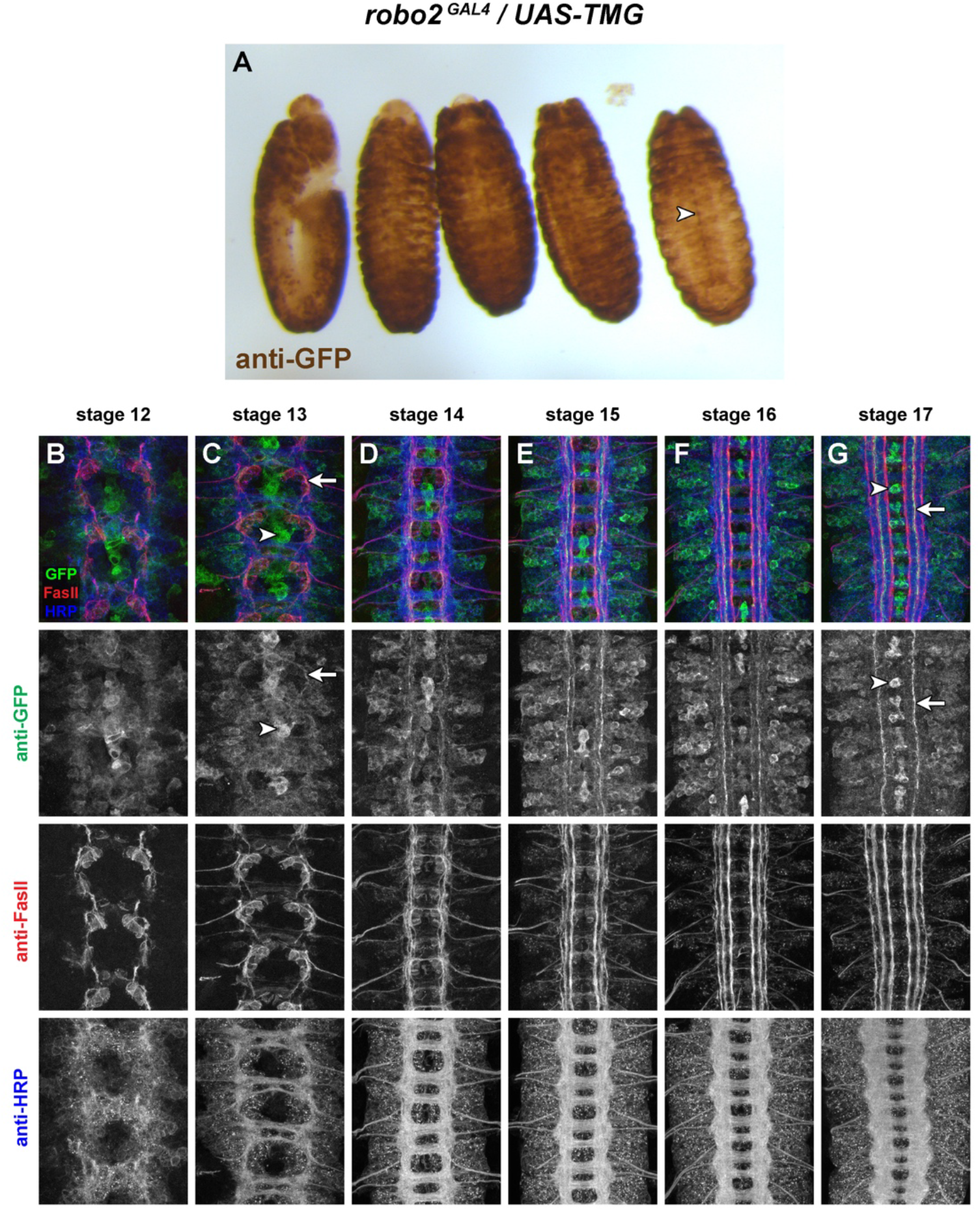
Expression of a *robo2^GAL4^* enhancer trap allele. (A) *robo2^GAL4^/UAS-TMG* embryos stained with an anti-GFP antibody (brown). Representative embryos from various developmental stages are arranged in order of increasing age from left to right. Lateral view of youngest embryo on left; all others ventral side up. GFP is broadly expressed throughout the epidermis in these embryos throughout embryonic development. GFP expression in midline cells of the ventral nerve cord is detectable (arrowhead) but other aspects of internal GFP expression are obscured by the strong epidermal staining. (B-G) Ventral nerve cords from stage 12-17 *robo2^GAL4^/UAS-TMG* embryos stained with anti-GFP (green), anti-FasII (red) and anti-HRP (blue) antibodies. Lower images show isolated channels for each of the three antibodies. GFP is strongly expressed in midline cells from stage 12-13 through stage 17 (C,G, arrowhead), and is detectable on early pioneer axons (C, arrow) that remain detectable in medial and intermediate longitudinal pathways in older embryos (G, arrow). As in the whole mount embryos in (A), the broad GFP expression throughout the nerve cord makes it difficult to identify other specific subsets of GFP-expressing neurons in these embryos.

Similarly, fluorescent staining with anti-GFP antibodies in combination with anti-HRP and anti-Fasll in *robo2^GAL4^/UAS-TMG* dissected embryonic ventral nerve cords reveals broad GFP expression at all stages of ventral nerve cord development **(Figure 3B-G).** The strongest expression is in midline glia and midline-adjacent pioneer neurons; by early stage 13 the axons of these neurons are clearly distinguishable and strong GFP expression remains in these cells through stage 17, where their axons projecting in intermediate longitudinal pathways remain strongly GFP-positive **(Figure 3G).** The lower-level broad GFP expression throughout the neuropile makes it difficult to clearly identify other specific subsets of GFP-positive axons in these embryos, or to connect GFP-positive axons with the corresponding neuronal cell bodies in order to identify specific subset(s) of *robo2-*expressing neurons.

Because the *P{GawB}NP6273* transgene is located near the *robo2* promoter, it is likely to be influenced by all of the enhancer elements that normally regulate *robo2* expression. Given that *robo2* is broadly and dynamically transcribed throughout the ventral nerve cord and other tissues at various developmental stages, and that the GAL4 and GFP proteins likely perdure in cells long after transcription ceases from the transgene’s promoter, it is not surprising that most cells that express *robo2* during embryonic development may remain GFP-positive through later stages. Examining the expression patterns conferred by isolated putative enhancer regions may allow a more precise characterization of *robo2* expression in specific subsets of neurons in addition to other non-neural embryonic tissues. We therefore now turn to a characterization of expression patterns for each of the 17 *robo2* cloned enhancer fragment transgenes.

### Summary of whole-embryo expression in each *robo2* GAL4 line

In the following section, we summarize the GFP expression patterns in embryos carrying each GAL4 transgene in combination with *UAS-TMG,* and describe specific aspects of ventral nerve cord expression where applicable. Whole-mount embryos and confocal z-stack projection images of dissected ventral nerve cords from embryonic developmental stages 12-17 are shown in a corresponding figure for each line **(Figures 4–20).**

**Figure 4.**
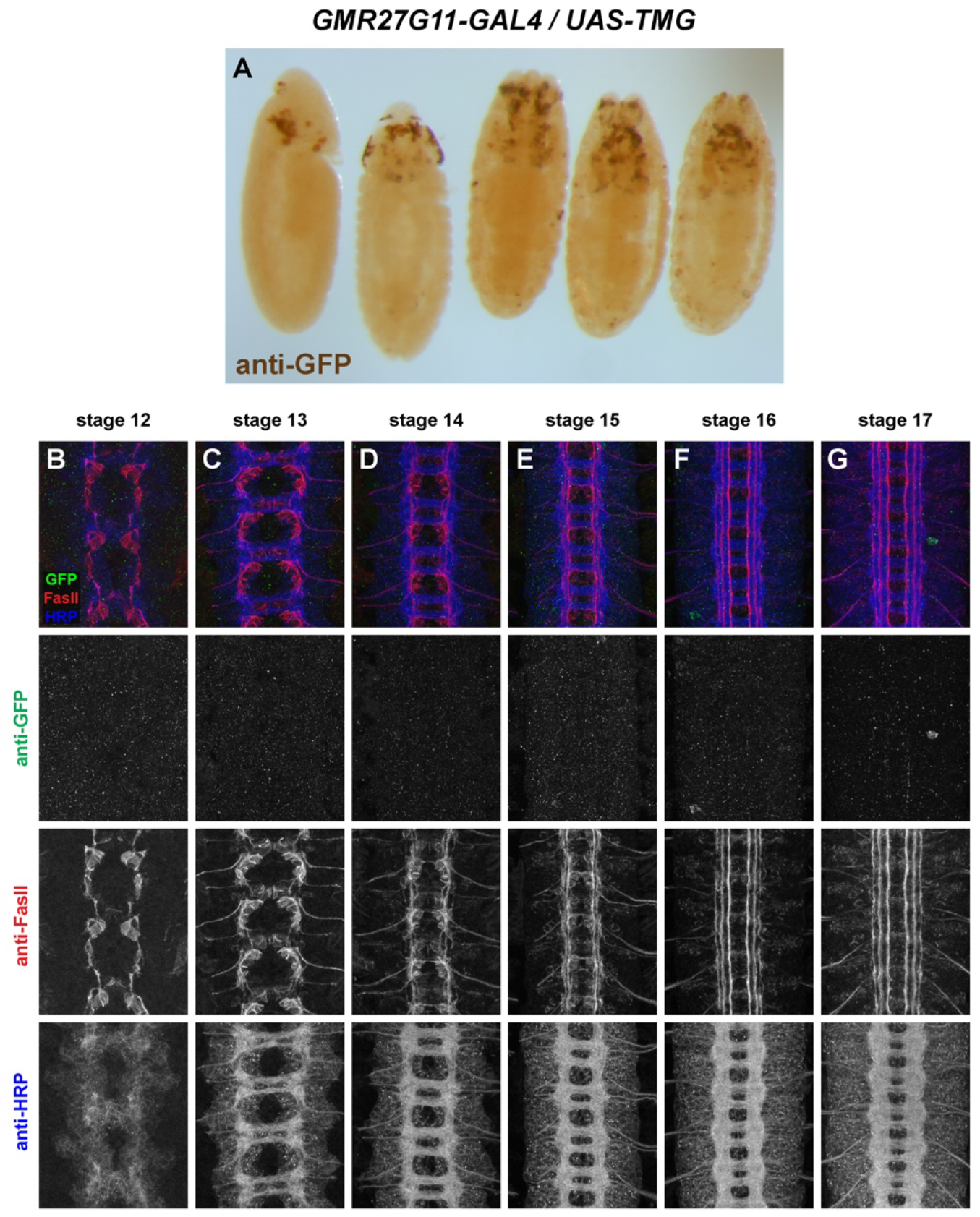
GMR27G11 lacks expression in the embryonic ventral nerve cord. (A) *GMR27G11/UAS-TMG* embryos stained with an anti-GFP antibody (brown). Representative embryos from various developmental stages are arranged in order of increasing age from left to right. Lateral view of youngest embryo on left; all others ventral side up. Little or no GFP expression is detectable in the ventral nerve cord. Sparse GFP expression is detectable in the anterior/head region in embryos from approximately stage 11 onwards. (B-G) Ventral nerve cords from stage 12-17 *GMR27G11/UAS-TMG* embryos stained with anti-GFP (green), anti-FasII (red) and anti-HRP (blue) antibodies. Lower images show isolated channels for each of the three antibodies. There is little or no GFP expression in the ventral nerve cord at any of the examined developmental stages.

#### *GMR27G11* (Figure 4)

No ventral nerve cord expression. Sparse expression in anterior/head begins around stage 11/12 and persists through stage 17.

#### *GMR27F07* (Figure 5)

Broadly expressed throughout brain and ventral nerve cord, beginning around stage 11/12 and persisting through stage 17. Expression in a few peripheral cells (possibly PNS neurons) begins around stage 13, with afferent processes visible reaching the nerve cord by stage 15. Expression in pioneer VNC neurons at stage 12-13, some FasII-positive and some FasII-negative. GFP-positive axons are present in medial and intermediate longitudinal pathways by stage 14, and in lateral longitudinal pathways by stage 15. GFP persists in all three longitudinal pathways through stage 17. Very little expression outside the nervous system.

**Figure 5.**
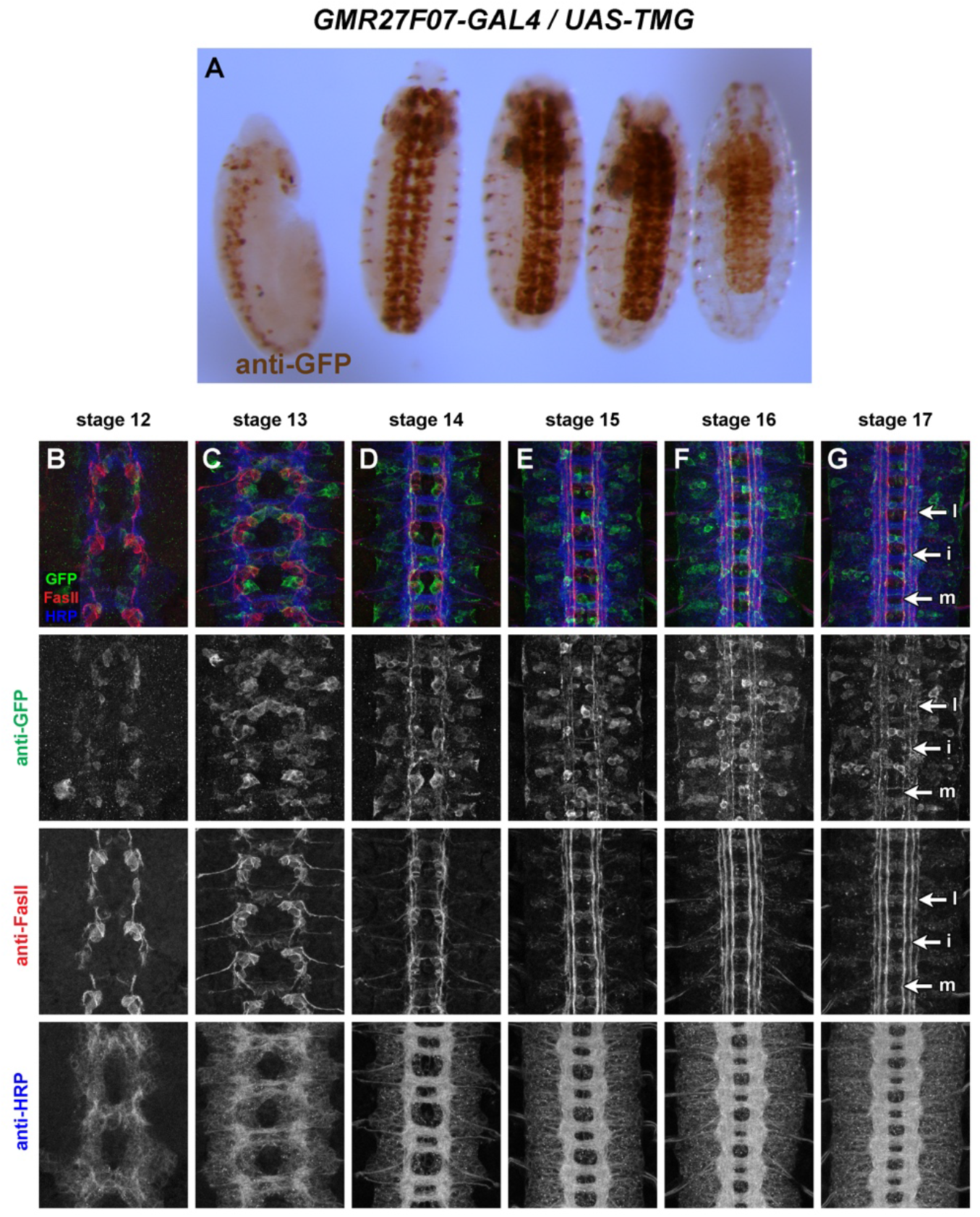
GMR27F07 is expressed broadly in the embryonic ventral nerve cord. (A) *GMR27F07/UAS-TMG* embryos stained with an anti-GFP antibody (brown). Representative embryos from various developmental stages are arranged in order of increasing age from left to right. Lateral view of youngest embryo on left; all others ventral side up. GFP is expressed broadly throughout the brain and ventral nerve cord beginning around stage 11/12, and is also detectable in peripheral nervous system cells beginning around stage 13. (B-G) Ventral nerve cords from stage 12-17 *GMR27F07/UAS-TMG* embryos stained with anti-GFP (green), anti-FasII (red) and anti-HRP (blue) antibodies. Lower images show isolated channels for each of the three antibodies. GFP is expressed in pioneer neurons as early as stage 12 (B) and broad expression persists through stage 17 (G). GFP-positive longitudinal axons are visible in medial, intermediate, and lateral pathways (G, arrows; “m”, medial; “i”, intermediate; “l”, lateral).

#### *GMR28D05* (Figure 6)

Strongly expressed throughout brain and ventral nerve cord, beginning around stage 11/12 and persisting through stage 17. Many commissural and longitudinal axons are labeled but broad expression makes it difficult to distinguish individual pathways or fasicles. Expression in a few peripheral cells (possibly PNS neurons) begins around stage 13. Likely peripheral glia exiting VNC are labeled at stage 13 and expression extends along perpipheral nerve persisting through stage 17. Moderate expression throughout the gut/visceral mesoderm beginning around stage 13 and persisting through stage 17.

**Figure 6.**
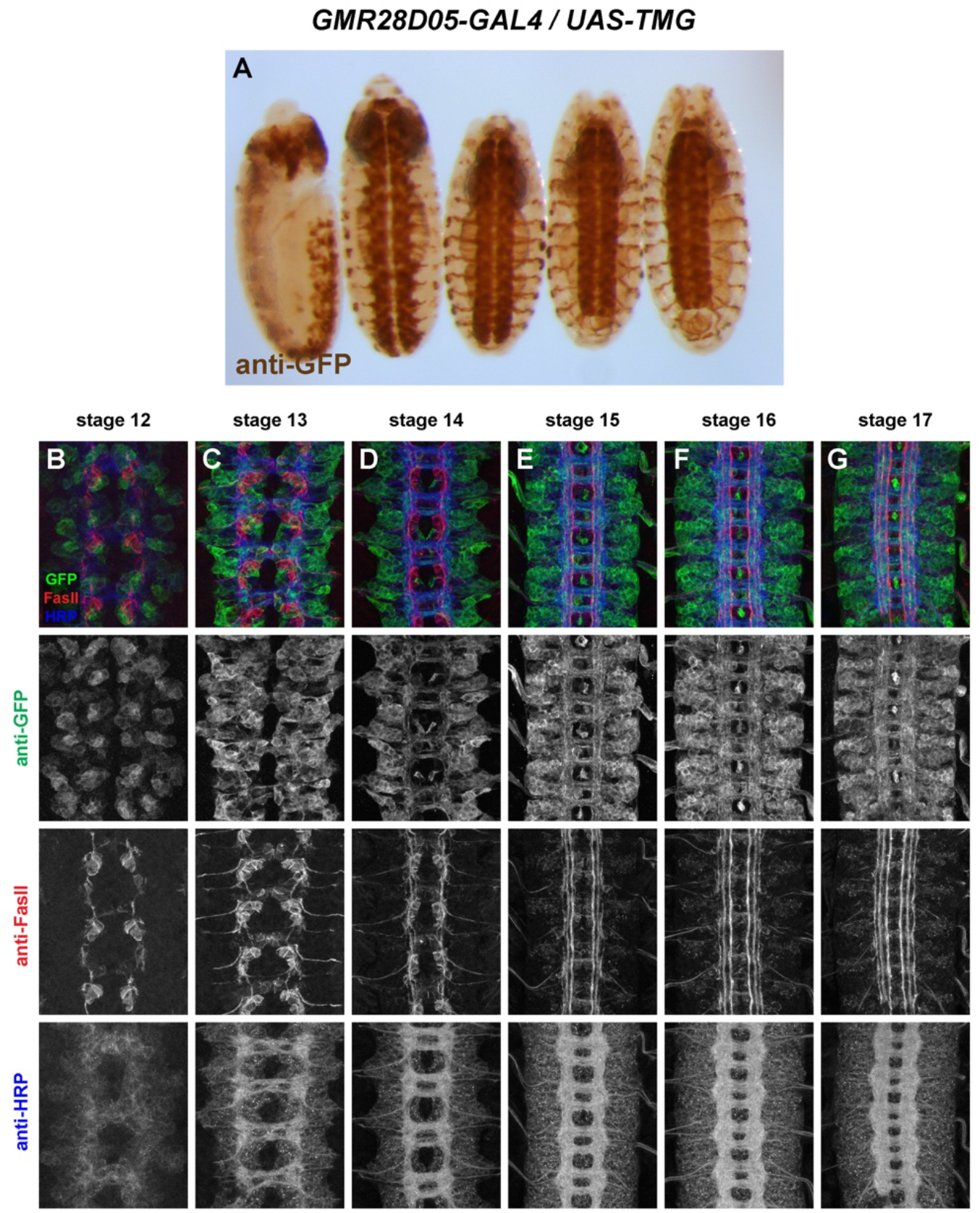
GMR28D05 is expressed broadly in the embryonic ventral nerve cord. (A) *GMR28D05/UAS-TMG* embryos stained with an anti-GFP antibody (brown). Representative embryos from various developmental stages are arranged in order of increasing age from left to right. Lateral view of youngest embryo on left; all others ventral side up. GFP is expressed broadly throughout the brain and ventral nerve cord beginning around stage 11/12, and is also detectable in peripheral nervous system cells beginning around stage 13. Moderate GFP expression throughout the gut begins around stage 13 and persists through stage 17. (B-G) Ventral nerve cords from stage 12-17 *GMR28D05/UAS-TMG* embryos stained with anti-GFP (green), anti-FasII (red) and anti-HRP (blue) antibodies. Lower images show isolated channels for each of the three antibodies. GFP is expressed broadly in VNC neurons as early as stage 12 (B) and broad expression persists through stage 17 (G). Many GFP-positive axons are visible in both commissures and many longitudinal pathways.

#### *GMR27D10* (Figure 7)

Broadly expressed in anterior/head and brain by stage 12, persisting through stage 17. Stochastic labeling of one or two ipsilateral pioneer neurons in some segments at stage 12/13 (likely vMP2/dMP2) whose axons remain in the medial FasII pathway through stage 17. Some expression in anterior head lobes at late stages (16/17). Little or no other expression outside of the CNS.

**Figure 7.**
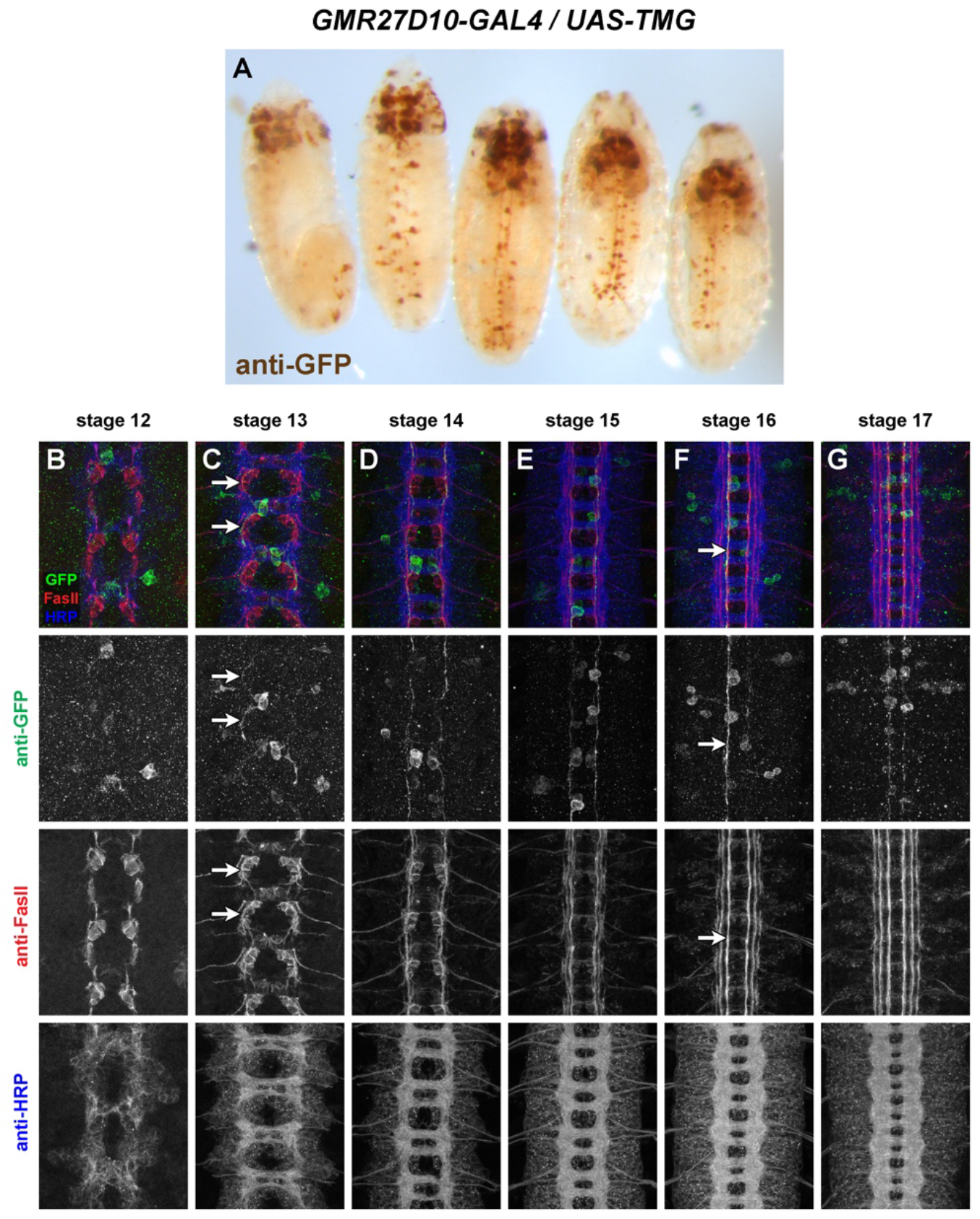
GMR27D10 is stochastically expressed in a small number of longitudinal pioneer neurons in the ventral nerve cord. (A) *GMR27D10/UAS-TMG* embryos stained with an anti-GFP antibody (brown). Representative embryos from various developmental stages are arranged in order of increasing age from left to right. Lateral view of youngest embryo on left; all others ventral side up. GFP is expressed in a few midline-adjacent ipsilateral neurons per hemisegment at stage 12-13, with scattered expression in a few other neurons farther from the midline. GFP persists in these neurons and their axons through stage 17. GFP expression in brain/head begins as early as stage 11 and persists through stage 17. (B-G) Ventral nerve cords from stage 12-17 *GMR27D10/UAS-TMG* embryos stained with anti-GFP (green), anti-FasII (red) and anti-HRP (blue) antibodies. Lower images show isolated channels for each of the three antibodies. GFP-positive pioneer neurons are likely dMP2/vMP2 based on cell body position near the midline, axon projection pattern (C, arrows; one projects dorsally and one ventrally), and the location of their axons in the medial longitudinal pathway at late stages (F, arrow). GFP expression is stochastic in these neurons and either zero, one, or two of them express GFP in each hemisegment.

#### *GMR27D07* (Figure 8)

Broadly expressed in anterior/head and brain by stage 12, persisting through stage 17. Expressed in clusters of neurons near the midline beginning around stage 13, including some commissural pioneer neurons. GFP persists in these neurons and their axons in or near medial and intermediate FasII pathways through stage 17. Expression in peripheral cells (possibly PNS neurons) begins around stage 13. Sensory and/or motor axons labeled beginning stage 14 through stage 17.

**Figure 8.**
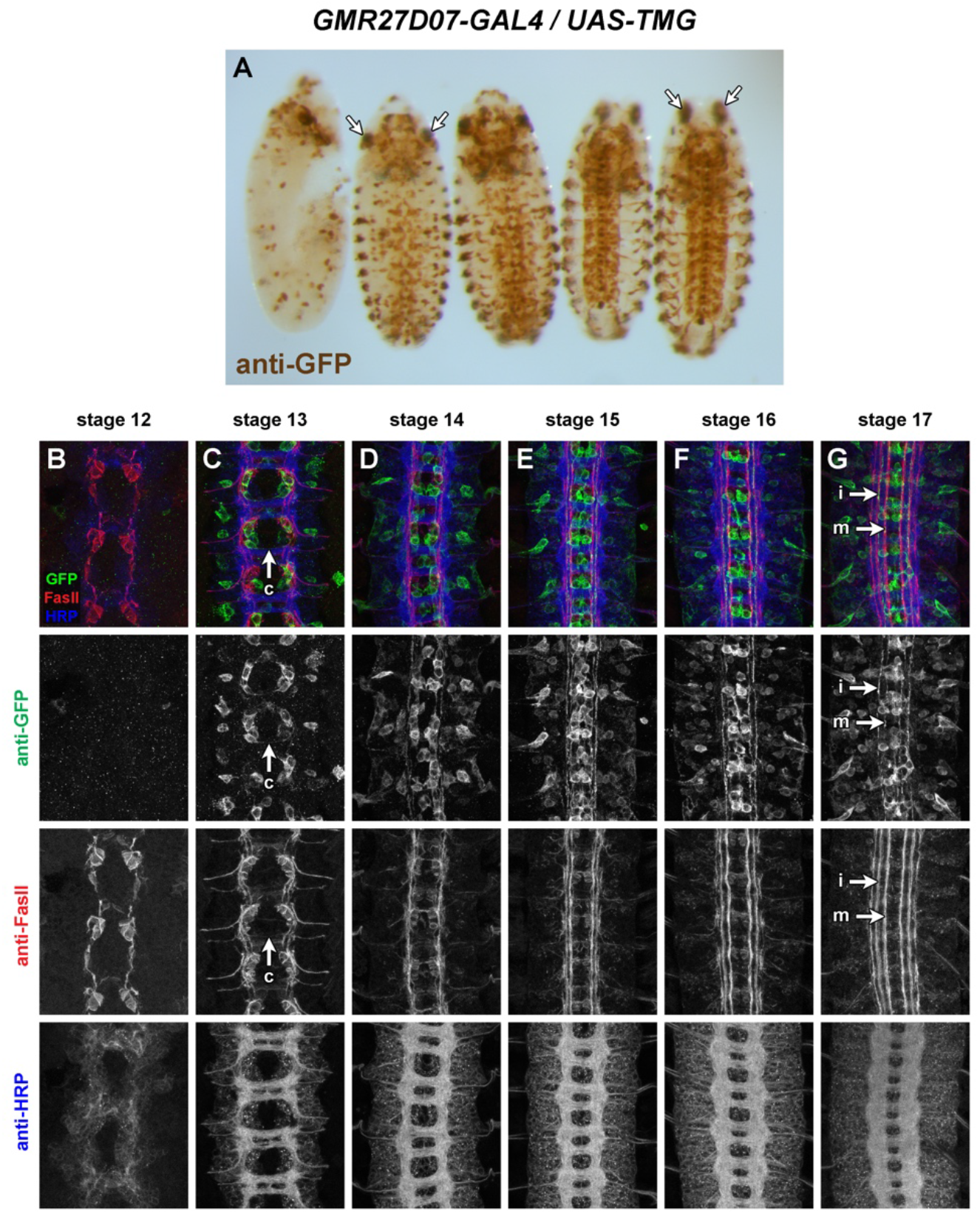
GMR27D07 is expressed in commissural pioneer and longitudinal neurons in the ventral nerve cord. (A) *GMR27D07/UAS-TMG* embryos stained with an anti-GFP antibody (brown). Representative embryos from various developmental stages are arranged in order of increasing age from left to right. Lateral view of youngest embryo on left; all others ventral side up. GFP is expressed in clusters of neurons in the ventral nerve cord and throughout the brain/head region beginning by stages 12-13 and persisting through stage 17. Strong midline staining and distinct longitudinal pathways visible through stage 17. Peripheral GFP staining begins around stage 13. Strong GFP expression in lateral spots in the head that migrate anteriorly in later embryos (arrows). (B-G) Ventral nerve cords from stage 12-17 *GMR27D07/UAS-TMG* embryos stained with anti-GFP (green), anti-FasII (red) and anti-HRP (blue) antibodies. Lower images show isolated channels for each of the three antibodies. Little or no GFP expression at stage 12; expression in small clusters of neurons begins stage 13, including some commissural pioneer neurons (C, arrow; “c”, commissural). Two distinct GFP-positive longitudinal pathways detectable by stage 14, persisting through stage 17 (G, arrows; “m”, medial; “i”, intermediate. Clusters of cells near the midline retain strong GFP expression through stage 17, with lower expression in a larger number of cells farther from the midline.

#### *GMR28A12* (Figure 9)

Little or no GFP expression is detectable in the ventral nerve cord, apart from a single bilateral pair of cells in each segment beginning around stage 13, located dorsally to the neuropile with processes extending laterally out of the CNS, likely dorsal channel glia (Ito *et al.* 1995) or transverse nerve (TN) exit glia (Gorczyca *et al.* 1994). Expressed in visceral mesoderm beginning stage 12 (two broad longitudinal stripes at stage 13/14, expanding to cover entire gut by stage 15). Strongly expressed in salivary glands beginning around stage 14/15. Expression in anterior/head beginning stage 12 or earlier, persisting through stage 17 including anterior head lobes.

**Figure 9.**
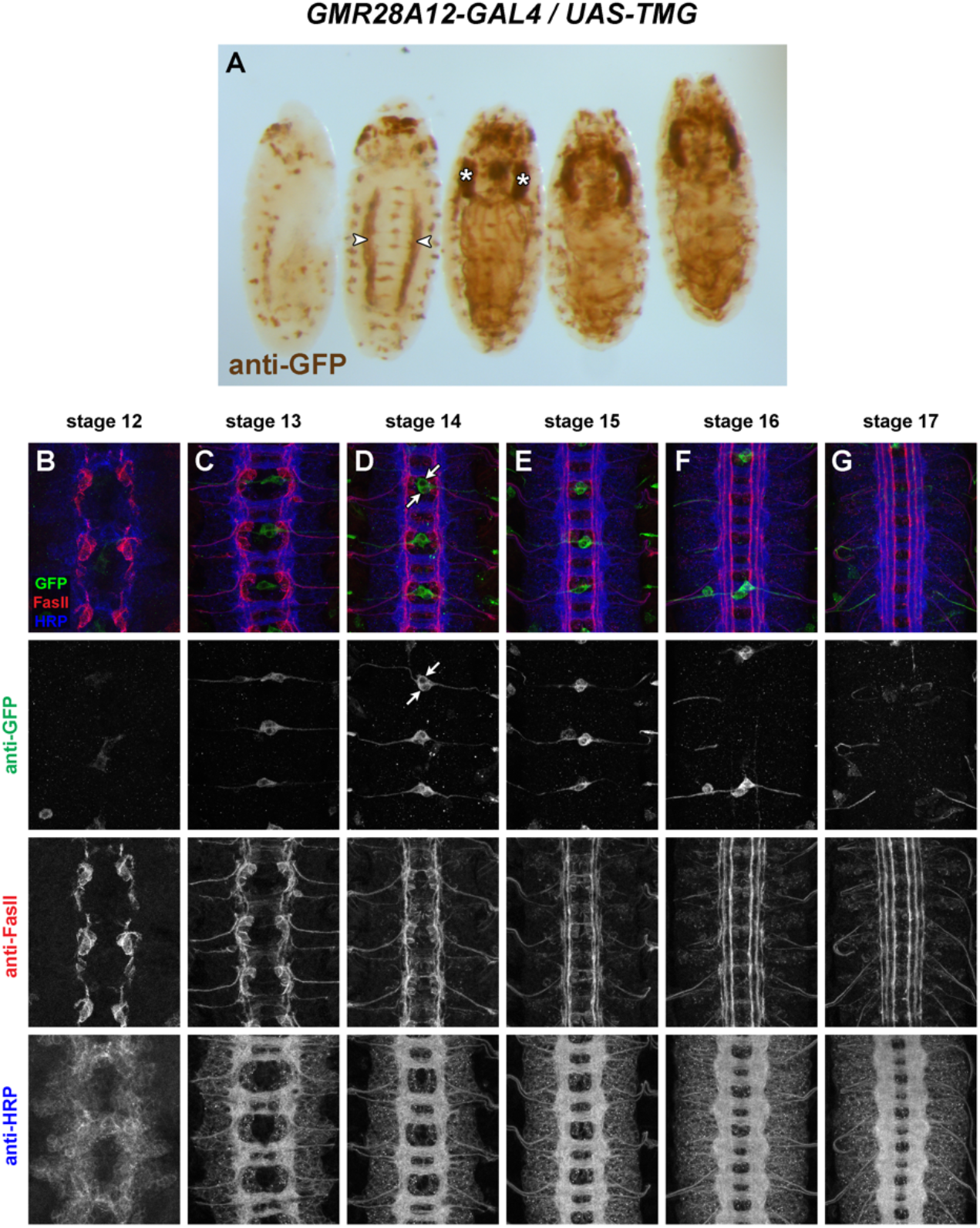
GMR28A12 exhibits little or no expression in the embryonic ventral nerve cord. (A) *GMR28A12/UAS-TMG* embryos stained with an anti-GFP antibody (brown). Representative embryos from various developmental stages are arranged in order of increasing age from left to right. Lateral view of youngest embryo on left; all others ventral side up. Little or no GFP expression is detectable in the ventral nerve cord, apart from a single bilateral pair of cells in each segment with processes extending laterally out of the CNS. GFP expressed in visceral mesoderm beginning stage 12 (arrowheads). GFP strongly expressed in salivary glands beginning around stage 14/15 (asterisks). Expression in anterior/head beginning stage 12 or earlier, persisting through stage 17 including anterior head lobes. (B-G) Ventral nerve cords from stage 12-17 *GMR28A12/UAS-TMG* embryos stained with anti-GFP (green), anti-FasII (red) and anti-HRP (blue) antibodies. Lower images show isolated channels for each of the three antibodies. There is little or no GFP expression in the ventral nerve cord at any of the examined developmental stages. Cells near the midline with lateral processes are located dorsally to the ventral nerve cord, likely dorsal channel glia/TN exit glia (D, arrows).

#### *GMR28F02* (Figure 10)

Expressed mainly in brain/VNC and visceral mesoderm. Visceral mesoderm expression begins by stage 12-13 (two broad longitudinal stripes at stage 13/14, expanding to cover entire gut by stage 15). In VNC, expressed consistently in a large cluster of neurons with lateral cell bodies plus some midline-adjacent neurons in each hemisegment beginning stage 12. Commissural axons from this cluster are detectable crossing the midline at stage 13: one bundle near posterior edge of PC and 2-3 bundles in the anterior half of AC. Lateral longitudinal axons are detectable at stage 13, forming an intact lateral pathway connecting adjacent segments by stage 14 (at this stage the lateral FasII pathways have not yet formed). Axon and cell body GFP persists through stage 17, where GFP+ longitudinal axons are detectable in or near the intermediate and lateral FasII pathways. Some motor nerve and/or peripheral glia expression visible by stage 15, persisting through stage 17. Expressed in dorsal channel glia/TN exit glia beginning around stage 13.

**Figure 10.**
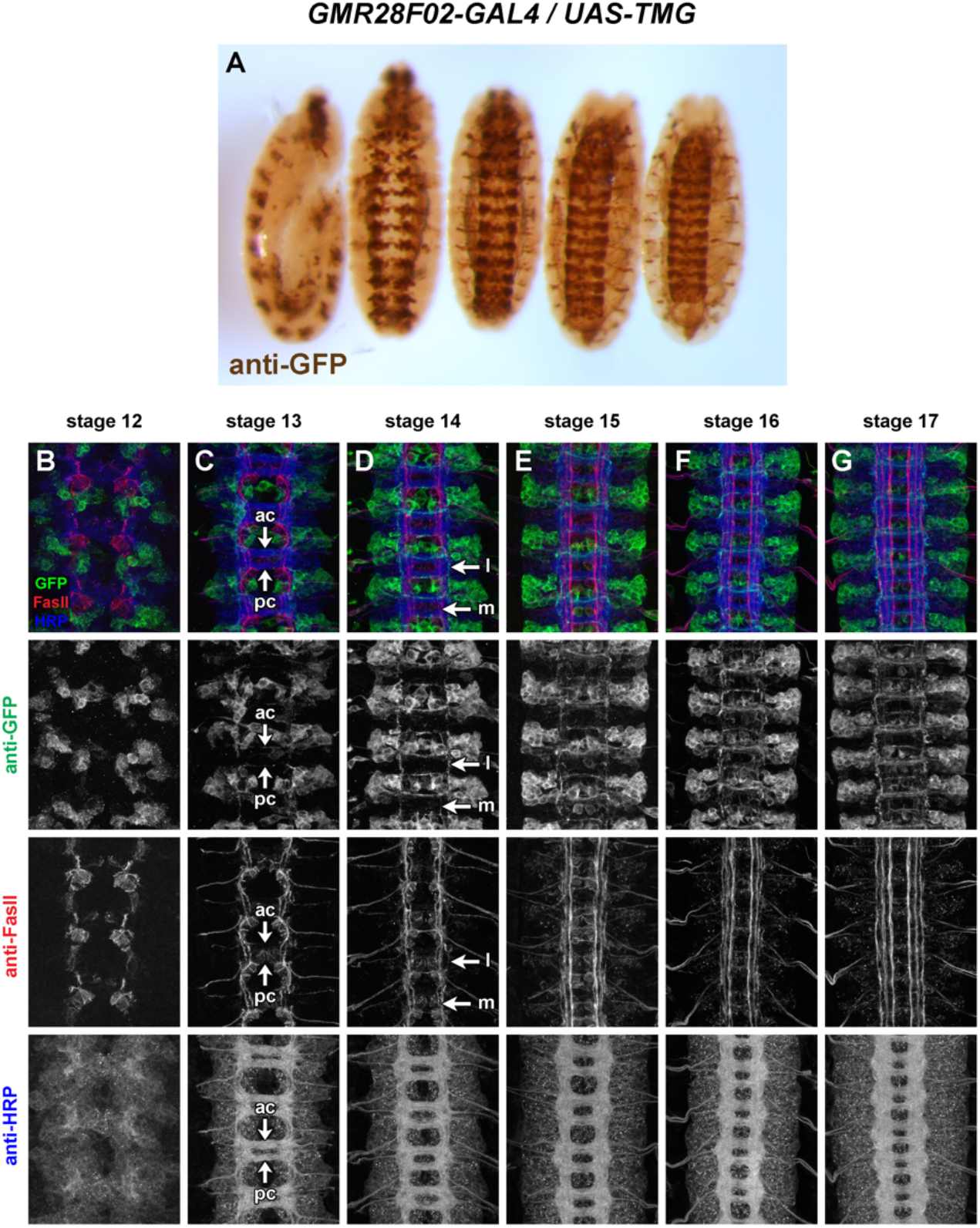
GMR28F02 is expressed in a subset of commissural longitudinal neurons. (A) *GMR28F02/UAS-TMG* embryos stained with an anti-GFP antibody (brown). Representative embryos from various developmental stages are arranged in order of increasing age from left to right. Lateral view of youngest embryo on left; all others ventral side up. Strong GFP expression in a segmentally repeated cluster of lateral neurons plus some midline-adjacent neurons. Peripheral GFP (possibly motor nerve or peripheral glia) detectable by stage 15. GFP detectable in visceral mesoderm from stage 12-13. Expressed in dorsal channel glia/TN exit glia beginning around stage 13. (B-G) Ventral nerve cords from stage 12-17 *GMR28F02/UAS-TMG* embryos stained with anti-GFP (green), anti-FasII (red) and anti-HRP (blue) antibodies. Lower images show isolated channels for each of the three antibodies. Medial and lateral neuronal cell bodies are GFP-positive by stage 12 (B). Commissural axons visible crossing the midline in both commissures by stage 13 (C, arrows; “ac”, anterior commissure; “pc”, posterior commissure), and turn longitudinally at medial/intermediate and lateral positions by stage 14 (D, arrows; “m”, medial/intermediate; “l”, lateral). GFP-positive lateral pathway forms before FasII-positive axons are detectable in the lateral region (compare GFP and FasII staining in panel D).

#### *GMR28E10* (Figure 11)

Very broadly expressed including nerve cord and epidermis beginning at stage 11 or earlier. Broad expression in neuronal cell bodies from stage 12 through stage 17; few axons individually discernable due to uniform expression in nerve cord. The expression pattern of GMR28E10 looks very similar to that of GMR28D10, with which it overlaps.

**Figure 11.**
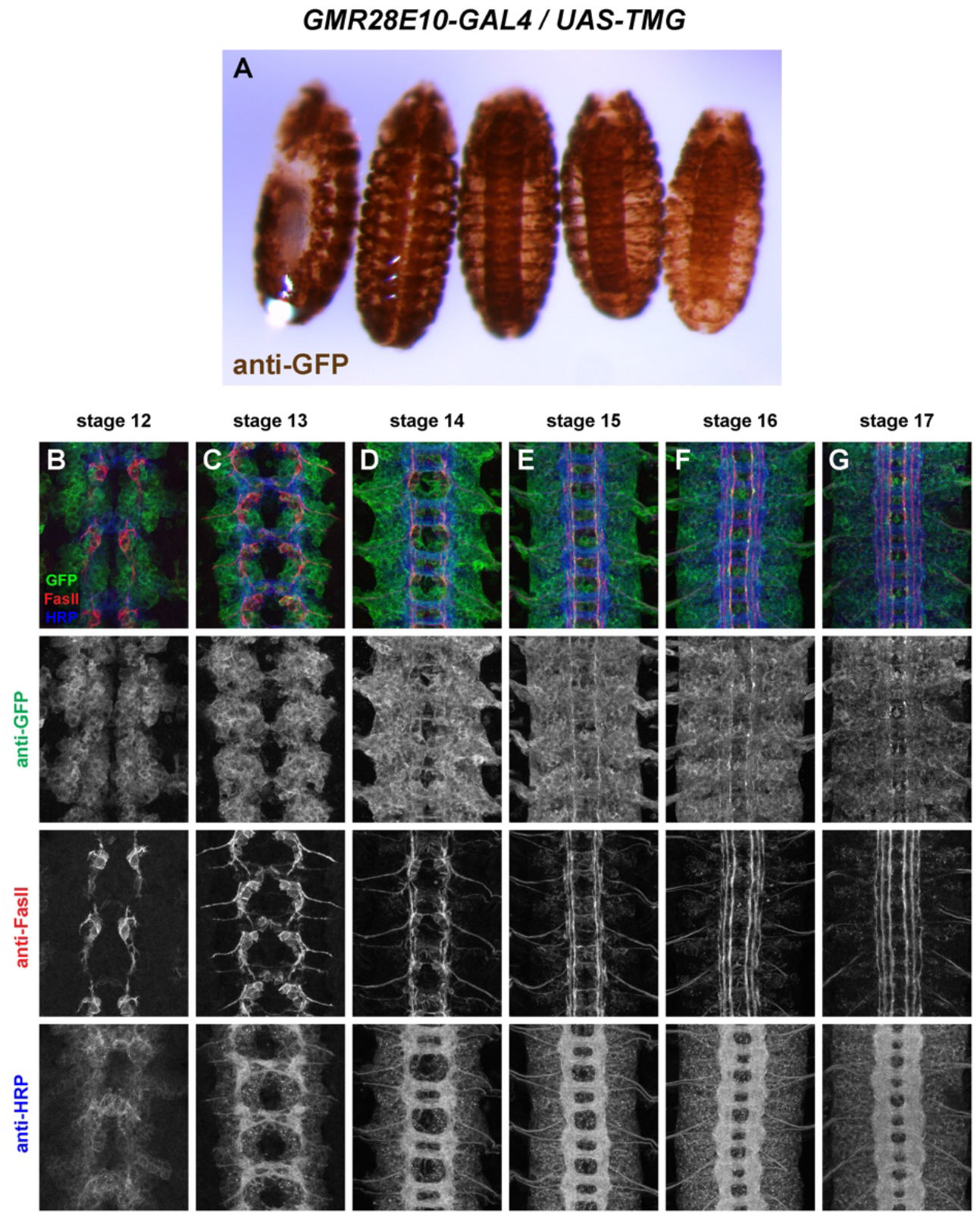
GMR28E10 is expressed broadly in the embryonic ventral nerve cord. (A) *GMR28E10/UAS-TMG* embryos stained with an anti-GFP antibody (brown). Representative embryos from various developmental stages are arranged in order of increasing age from left to right. Lateral view of youngest embryo on left; all others ventral side up. GFP is expressed broadly throughout the epidermis, brain, and ventral nerve cord beginning around stage 11/12. The broad staining makes it difficult to discern specific aspects of GFP expression in the ventral nerve cords of whole embryos. (B-G) Ventral nerve cords from stage 12-17 *GMR28E10/UAS-TMG* embryos stained with anti-GFP (green), anti-FasII (red) and anti-HRP (blue) antibodies. Lower images show isolated channels for each of the three antibodies. GFP is expressed broadly in many neurons from stage 12 through stage 17. While some distinct GFP-positive commissural and longitudinal axons are discernable, the broad GFP expression throughout the nerve cord makes it difficult to identify other specific subsets of GFP-expressing neurons in these embryos.

#### *GMR28D10* (Figure 12)

Very broadly expressed including nerve cord and epidermis beginning at stage 11 or earlier. Broad expression in neuronal cell bodies from stage 12 through stage 17; few axons individually discernable due to uniform expression in nerve cord. The expression pattern of GMR28D10 looks very similar to that of GMR28E10, with which it overlaps.

**Figure 12.**
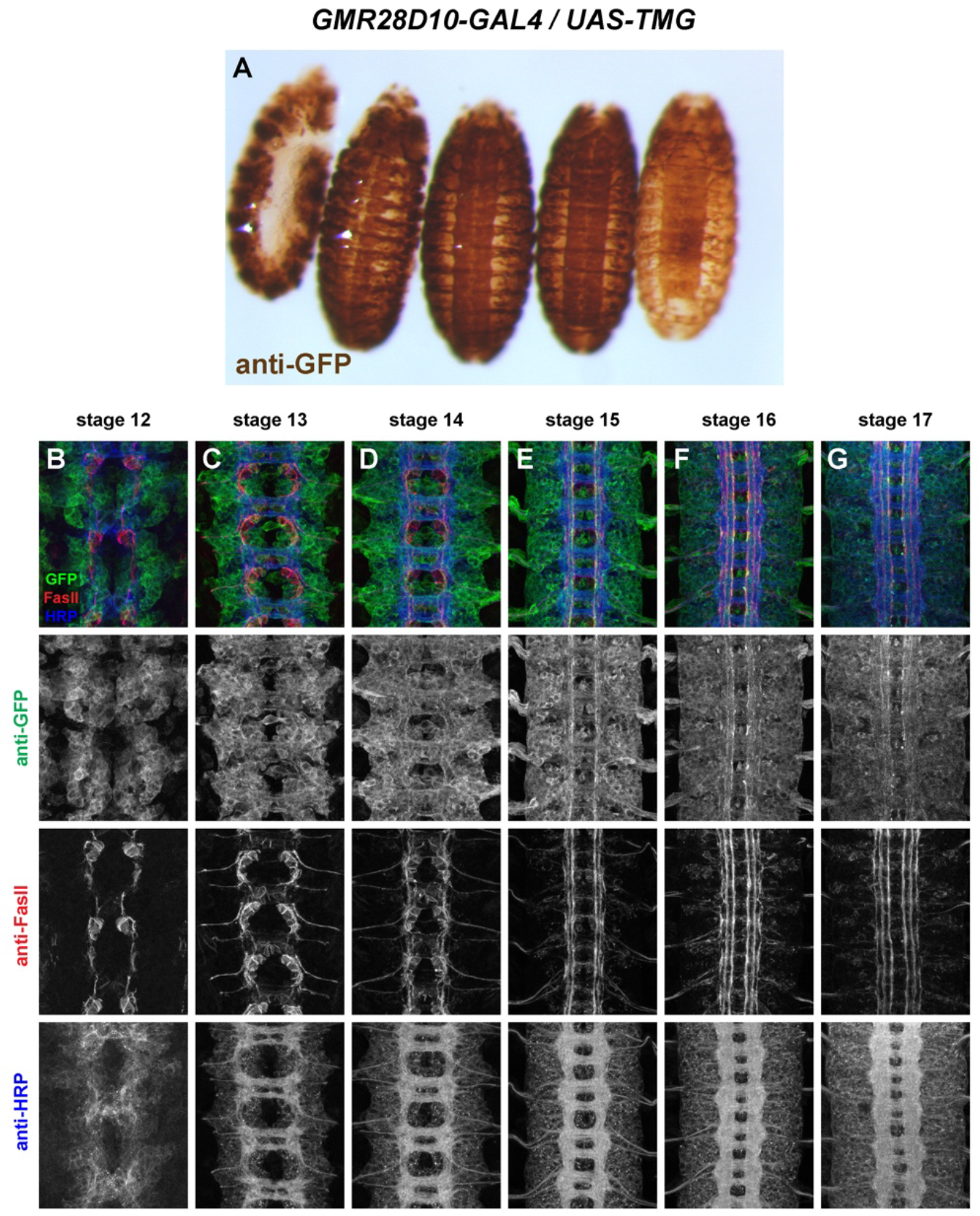
GMR28D10 is expressed broadly in the embryonic ventral nerve cord. (A) *GMR28D10/UAS-TMG* embryos stained with an anti-GFP antibody (brown). Representative embryos from various developmental stages are arranged in order of increasing age from left to right. Lateral view of youngest embryo on left; all others ventral side up. GFP is expressed broadly throughout the epidermis, brain, and ventral nerve cord beginning around stage 11/12. The broad staining makes it difficult to discern specific aspects of GFP expression in the ventral nerve cords of whole embryos. (B-G) Ventral nerve cords from stage 12-17 *GMR28D10/UAS-TMG* embryos stained with anti-GFP (green), anti-FasII (red) and anti-HRP (blue) antibodies. Lower images show isolated channels for each of the three antibodies. GFP is expressed broadly in many neurons from stage 12 through stage 17. While some distinct GFP-positive commissural and longitudinal axons are discernable, the broad GFP expression throughout the nerve cord makes it difficult to identify other specific subsets of GFP-expressing neurons in these embryos.

#### *GMR28G05* (Figure 13)

Expression mainly restricted to brain/head and ventral nerve cord. In VNC, expressed in a consistent cluster of neurons in each hemisegment with lateral cell bodies beginning stage 12. Commissural axons from this cluster are detectable crossing the midline at stage 13: one bundle near posterior edge of AC and 1-2 bundles at anterior edge of PC. Axons that cross in AC begin turning anteriorly by stage 14 and overlap with lateral FasII pathways by stage 15. Axons that cross in PC also appear to turn anteriorly by stage 14 but grow along medial or intermediate FasII pathways. Axon and cell body GFP persists through stage 17, where GFP+ longitudinal axons are detectable mainly in or near lateral FasII pathways, with a few in or near medial or intermediate FasII pathways. Peripheral glia expression is visible by stage 13/14 and persists through stage 17.

**Figure 13.**
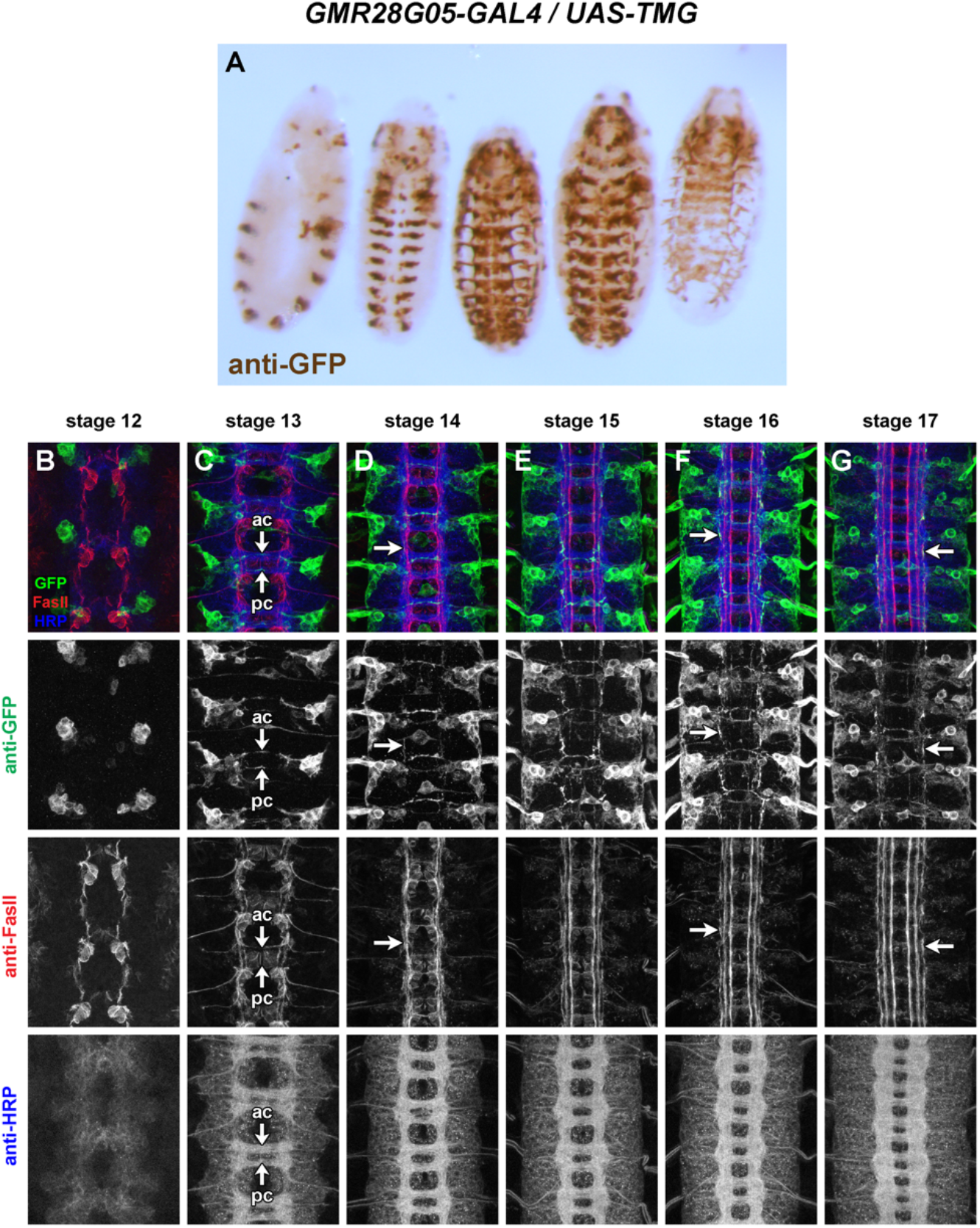
GMR28G05 is expressed in a subset of commissural longitudinal neurons. (A) *GMR28G05/UAS-TMG* embryos stained with an anti-GFP antibody (brown). Representative embryos from various developmental stages are arranged in order of increasing age from left to right. Lateral view of youngest embryo on left; all others ventral side up. Strong GFP expression in segmentally repeated clusters of neurons. GFP-positive cells wrapping external surface of nerve cord/motor nerve roots (likely peripheral glia) are prominent in mid-and late-stage embryos. (B-G) Ventral nerve cords from stage 12-17 *GMR28G05/UAS-TMG* embryos stained with anti-GFP (green), anti-FasII (red) and anti-HRP (blue) antibodies. Lower images show isolated channels for each of the three antibodies. Lateral neuronal cell bodies are GFP-positive by stage 12 (B). Commissural axons visible crossing the midline in both commissures by stage 13 (C, arrows; “ac”, anterior commissure; “pc”, posterior commissure), and turn longitudinally at medial/intermediate positions by stage 14 (D, arrow). GFP-positive axons are detectable in lateral pathways at stages 16-17, at the same time or later than FasII-positive lateral axon pathways form (F,G, arrows).

#### *GMR28D12* (Figure 14)

Uniformly expressed throughout ectoderm beginning at stage 11 or earlier. Also uniformly expressed throughout VNC from stage 12 through 17. Higher expression in midline glia beginning at stage 12, persisting through stage 17. Similar expression to GMR28B05 except lower in midline glia.

**Figure 14.**
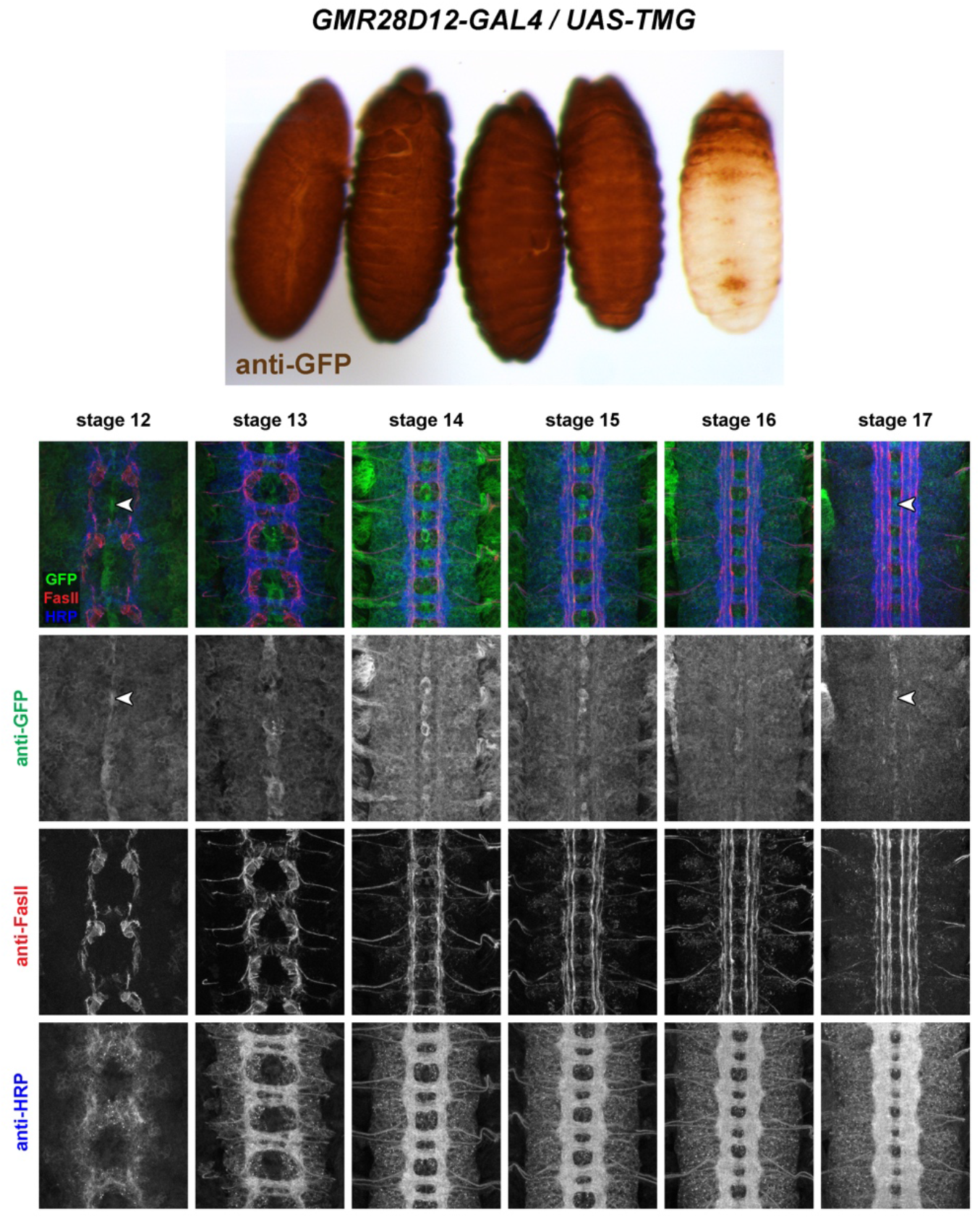
GMR28D12 is expressed broadly in the embryonic ventral nerve cord. (A) *GMR28D12/UAS-TMG* embryos stained with an anti-GFP antibody (brown). Representative embryos from various developmental stages are arranged in order of increasing age from left to right. Lateral view of youngest embryo on left; all others ventral side up. GFP is expressed broadly throughout the epidermis. The broad staining makes it difficult to discern specific aspects of GFP expression in the ventral nerve cords of whole embryos. (B-G) Ventral nerve cords from stage 12-17 *GMR28D12/UAS-TMG* embryos stained with anti-GFP (green), anti-FasII (red) and anti-HRP (blue) antibodies. Lower images show isolated channels for each of the three antibodies. GFP is expressed broadly in many neurons and elevated in midline cells from stage 12 (B, arrowhead) through stage 17 (G, arrowhead). The broad GFP expression makes it difficult to identify specific subsets of GFP-expressing neurons in these embryos.

#### *GMR28F12* (Figure 15)

No detectable VNC expression. Strong staining in some cells in brain/head region beginning around stage 13 and persisting through stage 17 in anterior head lobes.High expression in visceral mesoderm beginning around stage 13 (two broad longitudinal stripes at stage 13/14, expanding to cover entire gut by stage 15). Striped expression in ectoderm also beginning around stage 13. The visceral mesoderm and striped ectodermal expression in this line resembles the Robo2 expression pattern described by Kraut and Zinn (Kraut and Zinn 2004).

**Figure 15.**
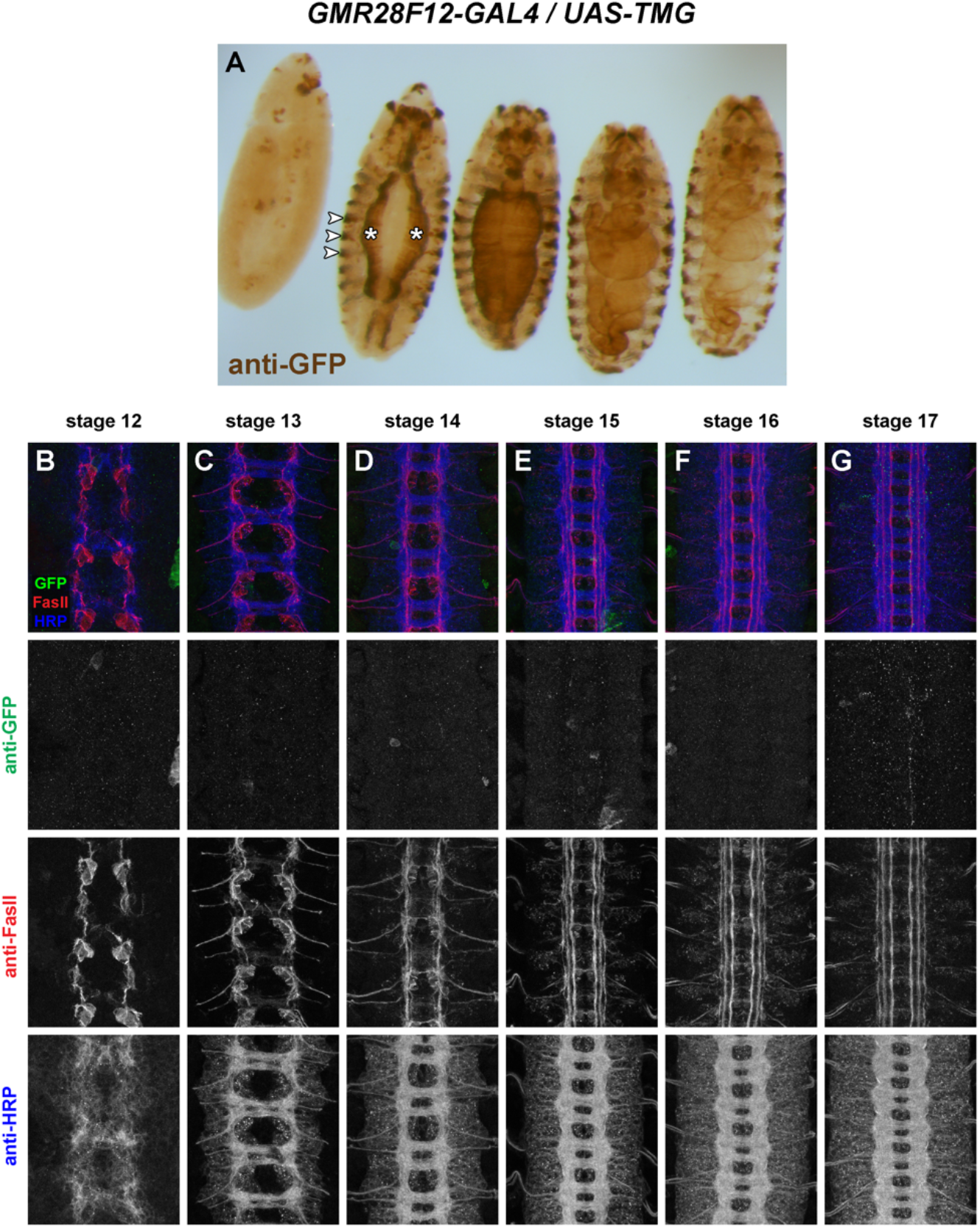
GMR28F12 lacks expression in the embryonic ventral nerve cord. (A) *GMR28F12/UAS-TMG* embryos stained with an anti-GFP antibody (brown). Representative embryos from various developmental stages are arranged in order of increasing age from left to right. Lateral view of youngest embryo on left; all others ventral side up. GFP expression is undetectable in the ventral nerve cord. GFP is strongly expressed in the visceral mesoderm (asterisks) and in ectodermal stripes (arrowheads) beginning around stage 13. Strong staining in some cells in brain/head region beginning around stage 13 and persisting through stage 17 in anterior head lobes. (B-G) Ventral nerve cords from stage 12-17 *GMR28F12/UAS-TMG* embryos stained with anti-GFP (green), anti-FasII (red) and anti-HRP (blue) antibodies. Lower images show isolated channels for each of the three antibodies. There is little or no GFP expression in the ventral nerve cord at any of the examined developmental stages.

#### *GMR27H02* (Figure 16)

Sparse but regular expression in a few neurons per hemisegment in the VNC beginning around stage 14, with lateral cell bodies and medial/intermediate ipsilateral axons. Expressed in dorsal channel glia/TN exit glia beginning around stage 13. Restricted expression in brain/head from stage 12 through stage 17, including anterior head lobes.

**Figure 16.**
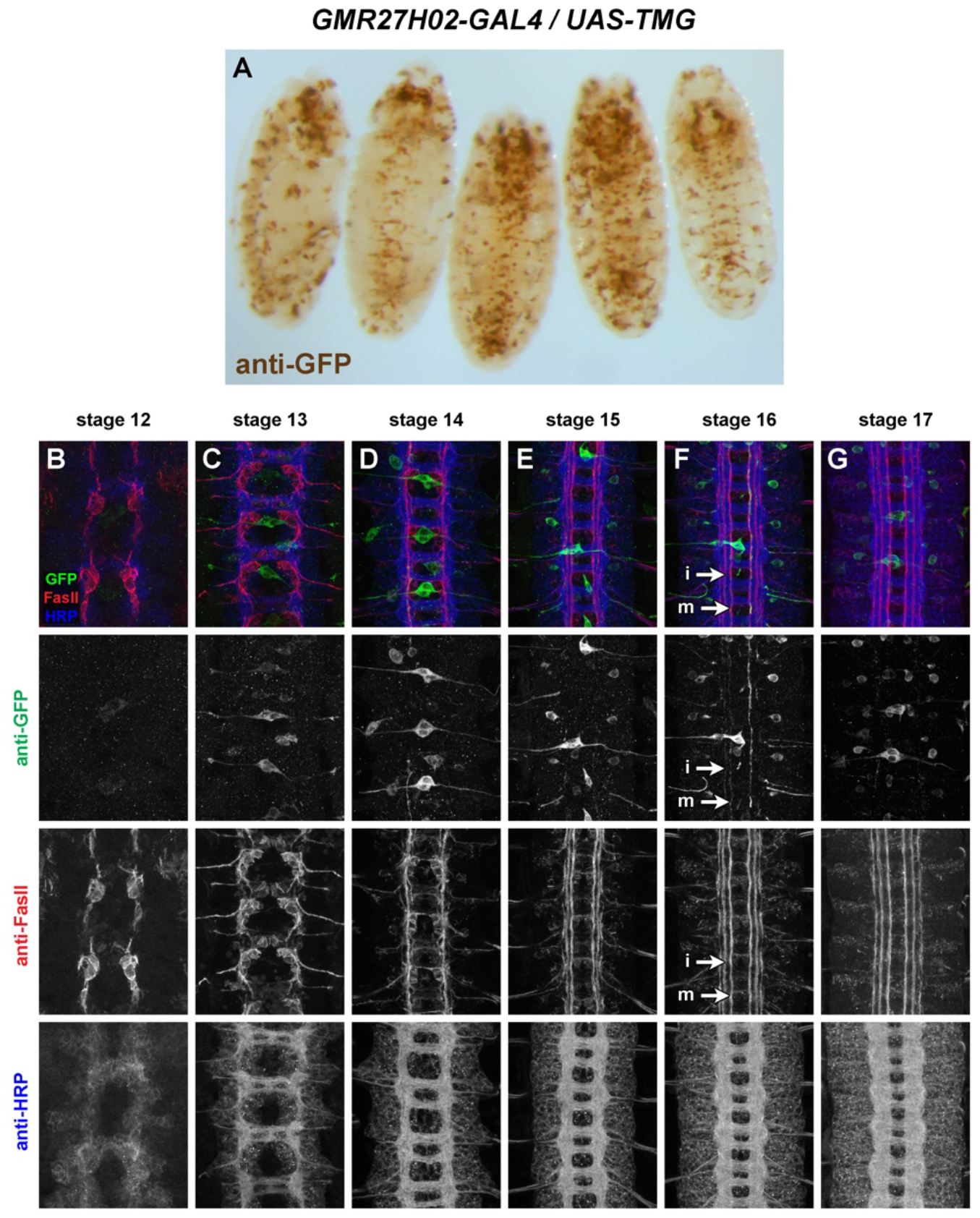
GMR27H02 is expressed in a small number of longitudinal neurons in the embryonic ventral nerve cord. (A) *GMR27H02/UAS-TMG* embryos stained with an anti-GFP antibody (brown). Representative embryos from various developmental stages are arranged in order of increasing age from left to right. Lateral view of youngest embryo on left; all others ventral side up. GFP is expressed throughout the head/brain beginning around stage 12, and sparsely expressed in a few individually identifiable neurons per segment in the VNC, along with dorsal channel glia/TN exit glia. (B-G) Ventral nerve cords from stage 12-17 *GMR27H02/UAS-TMG* embryos stained with anti-GFP (green), anti-FasII (red) and anti-HRP (blue) antibodies. Lower images show isolated channels for each of the three antibodies. The GFP-positive neurons are ipsilateral longitudinal neurons, with axons detectable in medial and intermediate axon pathways (F, arrows; “m”, medial; “i”, intermediate). Only two or three GFP-positive neurons are detectable in most abdominal VNC hemisegments.

#### *GMR28C04* (Figure 17)

Expression mainly restricted to brain/VNC and body wall muscles, in particular ventral muscles. Broad expression throughout VNC neurons starting at stage 13. Some commissural axons visible early (stage 13) and medial/intermediate longitudinal axons prominently labeled by stage 16/17. Two strong foci of expression in lateral head at stage 12, persist through stage 17 at tips of anterior head lobes.

**Figure 17.**
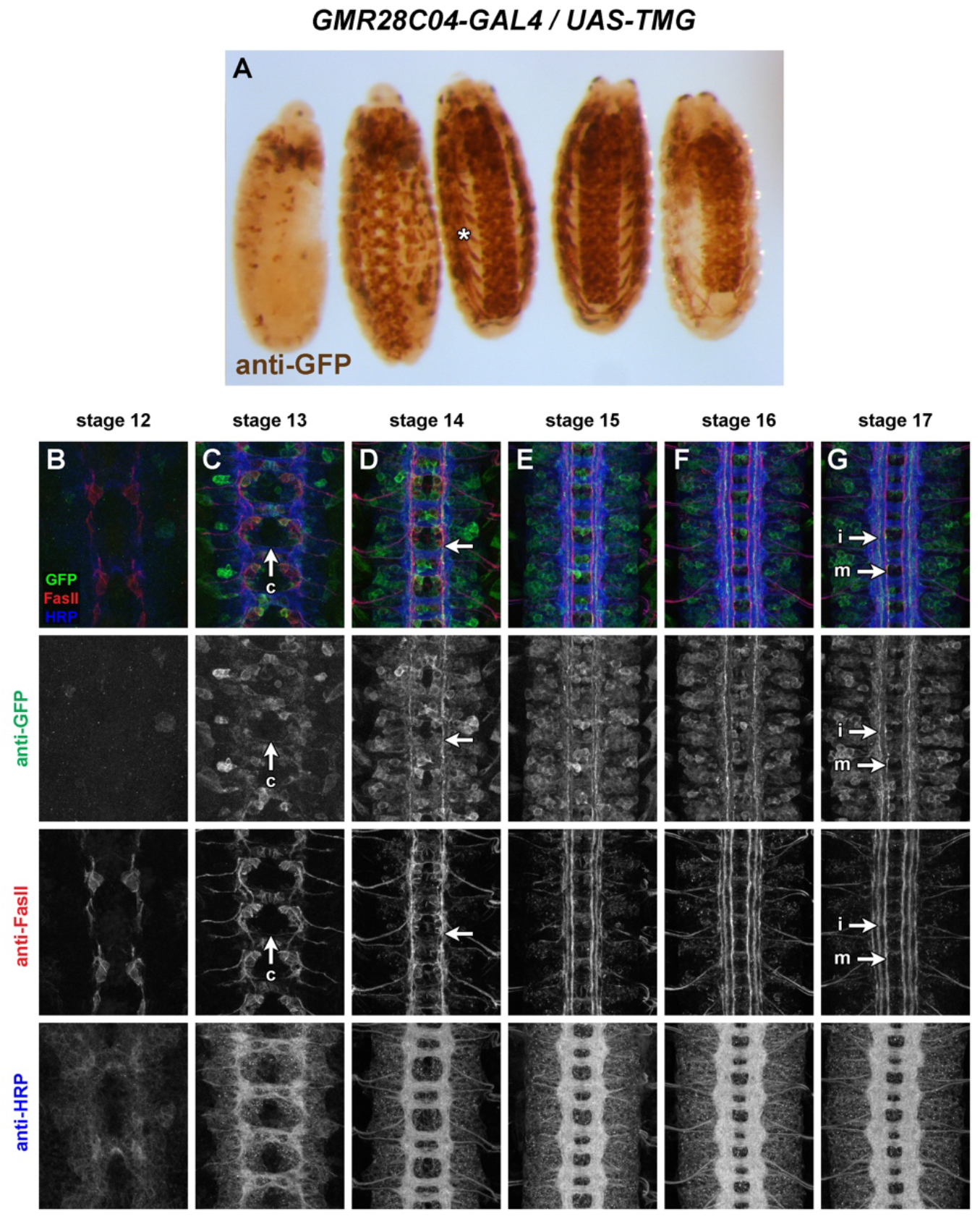
GMR28C04 is expressed broadly in the embryonic ventral nerve cord. (A) *GMR28C04/UAS-TMG* embryos stained with an anti-GFP antibody (brown). Representative embryos from various developmental stages are arranged in order of increasing age from left to right. Lateral view of youngest embryo on left; all others ventral side up. GFP is expressed broadly throughout the brain and ventral nerve cord beginning around stage 11/12. The broad staining makes it difficult to discern specific aspects of GFP expression in the ventral nerve cords of whole embryos. GFP is also broadly expressed in body wall muscles (A, asterisk). (B-G) Ventral nerve cords from stage 12-17 *GMR28C04/UAS-TMG* embryos stained with anti-GFP (green), anti-FasII (red) and anti-HRP (blue) antibodies. Lower images show isolated channels for each of the three antibodies. GFP is expressed broadly in many neurons beginning at stage 13. GFP-positive commissural axons are visible at stage 13 (C, arrow; “c”, commissural) and GFP-positive longitudinal axons are detectable beginning at stage 14 (D, arrow) and persisting through stage 17 in medial and intermediate positions (G, arrows; “m”, medial; “i”, intermediate). The broad GFP expression in the VNC makes it difficult to link these longitudinal axons to specific neuronal cell bodies.

#### *GMR28B05* (Figure 18)

Uniformly expressed throughout ectoderm beginning at stage 11 or earlier. Also uniformly expressed throughout VNC from stage 12 through 17. Higher expression in midline glia beginning at stage 12, persisting through stage 17. Similar expression to GMR28D12 except higher in midline glia.

**Figure 18.**
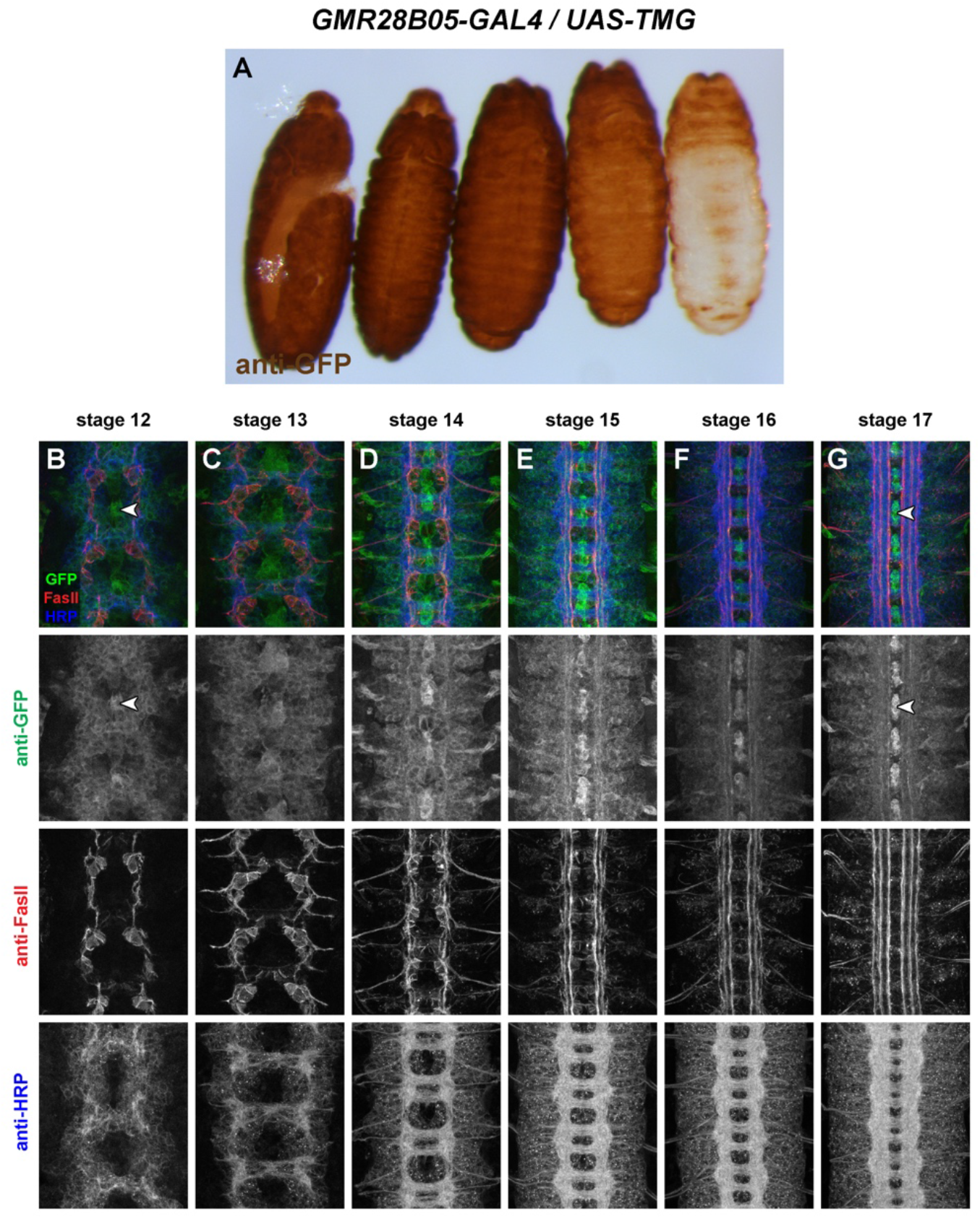
GMR28B05 is expressed broadly in the embryonic ventral nerve cord. (A) *GMR28B05/UAS-TMG* embryos stained with an anti-GFP antibody (brown). Representative embryos from various developmental stages are arranged in order of increasing age from left to right. Lateral view of youngest embryo on left; all others ventral side up. GFP is expressed broadly throughout the epidermis. The broad staining makes it difficult to discern specific aspects of GFP expression in the ventral nerve cords of whole embryos. (B-G) Ventral nerve cords from stage 12-17 *GMR28B05/UAS-TMG* embryos stained with anti-GFP (green), anti-FasII (red) and anti-HRP (blue) antibodies. Lower images show isolated channels for each of the three antibodies. GFP is expressed broadly in many neurons and especially high in midline cells from stage 12 (B, arrowhead) through stage 17 (G, arrowhead). The broad GFP expression makes it difficult to identify specific subsets of GFP-expressing neurons in these embryos, although some GFP-positive longitudinal axons in medial and intermediate pathways are detectable.

#### *GMR28A10* (Figure 19)

Little or no VNC expression. Head/anterior expression from stage 12 through stage 17, and scattered peripheral cells, possibly PNS neurons or peripheral glia.

**Figure 19.**
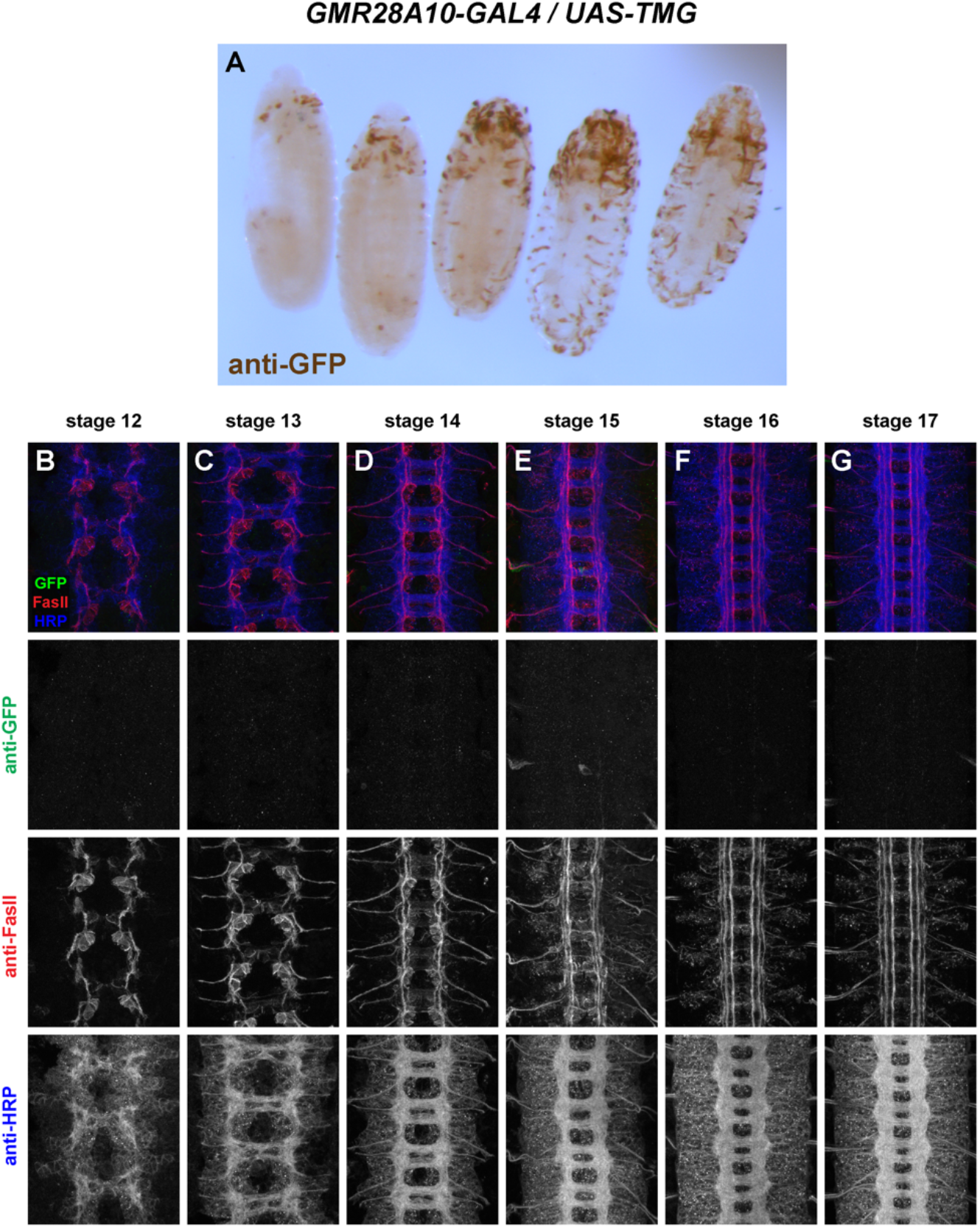
GMR28A10 exhibits little or no expression in the embryonic ventral nerve cord. (A) *GMR28A10/UAS-TMG* embryos stained with an anti-GFP antibody (brown). Representative embryos from various developmental stages are arranged in order of increasing age from left to right. Lateral view of youngest embryo on left; all others ventral side up. Little or no GFP expression is detectable in the ventral nerve cord. Scattered GFP expression in the anterior/head and in some peripheral cells at later stages. (B-G) Ventral nerve cords from stage 12-17 *GMR28A10/UAS-TMG* embryos stained with anti-GFP (green), anti-FasII (red) and anti-HRP (blue) antibodies. Lower images show isolated channels for each of the three antibodies. There is little or no GFP expression in the ventral nerve cord at any of the examined developmental stages.

#### *GMR28E07* (Figure 20)

Broadly expressed in nerve cord, body wall muscles/peripheral ectoderm, and head, beginning around stage 12/13 and persisting through stage 17. Strong expression in pioneer neurons near the midline and in midline glia at stage 12/13. Broad low-level expression throughout VNC persists through stage 17, along with higher expression in midline glia and midline-adjacent neurons with axons in medial and intermediate longitudinal pathways. Expressed in dorsal channel glia/TN exit glia beginning around stage 13.

**Figure 20.**
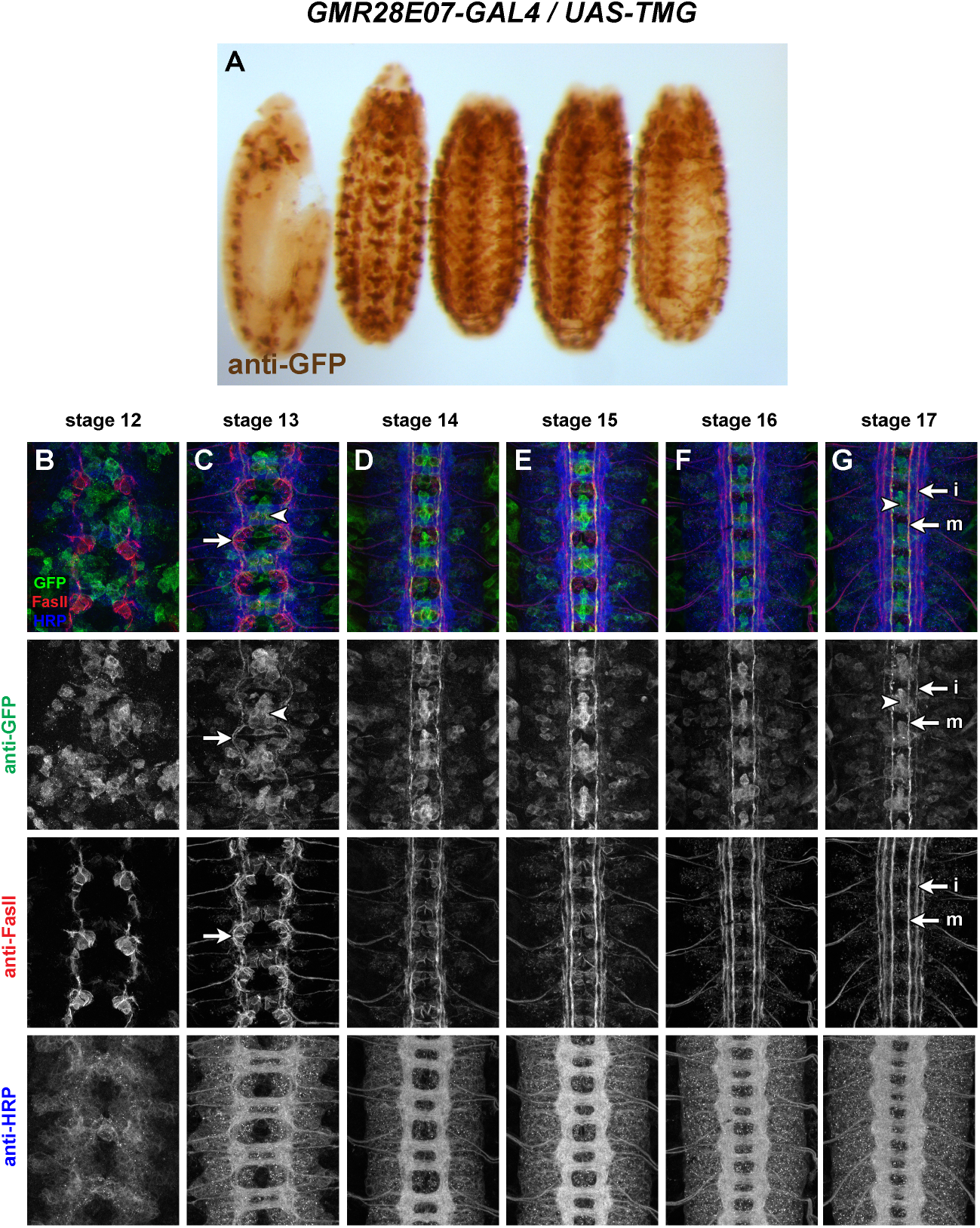
GMR28E07 is expressed strongly in midline glia and pioneer longitudinal neurons. (A) *GMR28E07/UAS-TMG* embryos stained with an anti-GFP antibody (brown). Representative embryos from various developmental stages are arranged in order of increasing age from left to right. Lateral view of youngest embryo on left; all others ventral side up. GFP is broadly expressed at a low level throughout the ventral nerve cord, and at a higher level in distinct cell clusters near the midline in each segment. GFP also expressed throughout the anterior/head and in body wall muscles and/or peripheral cells in the ectoderm. (B-G) Ventral nerve cords from stage 12-17 *GMR28E07/UAS-TMG* embryos stained with anti-GFP (green), anti-FasII (red) and anti-HRP (blue) antibodies. Lower images show isolated channels for each of the three antibodies. GFP is expressed in many neurons by stage 12, with little or no overlap with FasII (B). By stage 13 higher GFP expression is seen in midline cell clusters (C, arrowhead), with GFP-positive axons forming longitudinal tracts connecting adjacent segments (C, arrow; these tracts appear to be both GFP-positive and FasII-positive). GFP expression remains strong in midline cells (G, arrowhead) and longitudinal axons within medial and intermediate pathways (G, arrows; “m”, medial; “i”, intermediate) through stage 17. Expressed in dorsal channel glia/TN exit glia beginning around stage 13.

#### Enhancer fragments promoting *robo2* expression in midline glia

We have previously shown that *robo2* mRNA is transiently expressed in midline glia during the early stages of axon pathfinding (stages 12-13), and that the *robo2-GAL4* enhancer-trap insertion *P{GawB}NP6273* and HA-tagged Robo2 protein expressed from the *robo2^robo2^* knockin allele are also expressed in midline glia at these stages (Evans *et al.* 2015). Robo2 expression in midline cells is important for its pro-midline crossing activity, as it antagonizes Slit-Robo1 repulsion in trans in pre-crossing commissural axons to allow Robo1-expressing growth cones to cross the Slit-expressing midline (Evans *et al.* 2015).

Three of the lines described here are strongly expressed in midline cells, including cells with positions and morphologies consistent with midline glia (GMR28D12, GMR28B05, and GMR28E07) **(Figures 14, 18, 20)**. The GMR28E07 and GMR28B05 fragments are both located upstream of the *robo2* promoter but do not overlap with each other, while GMR28D12 is located within the first intron of *robo2.* Overall, GMR28E07 appears to most closely reproduce the midline expression seen in the *robo2-GAL4* line, with strong GFP expression in midline cells (both midline glia and midline-adjacent neurons) beginning early (stage 12-13) and persisting through stage 17 (compare **Figure 3** with **Figure 20**). Both *robo2-GAL4/UAS-TMG* and *GMR28E07/UAS-TMG* embryos also include GFP-labeled neuronal axons visible in the medial (pCC/dMP2) pathway and intermediate (MP1) pathway. To confirm midline glial expression of GMR28E07, we stained *GMR28E07/UAS-TMG* embryos with anti-wrapper, a marker for midline glia (Noordermeer *et al.* 1998), along with anti-GFP. We found that GFP clearly labeled wrapper-positive midline glia at stage 13, and GFP/wrapper co-localization persisted through stage 17 **(Figure 21).**

**Figure 21.**
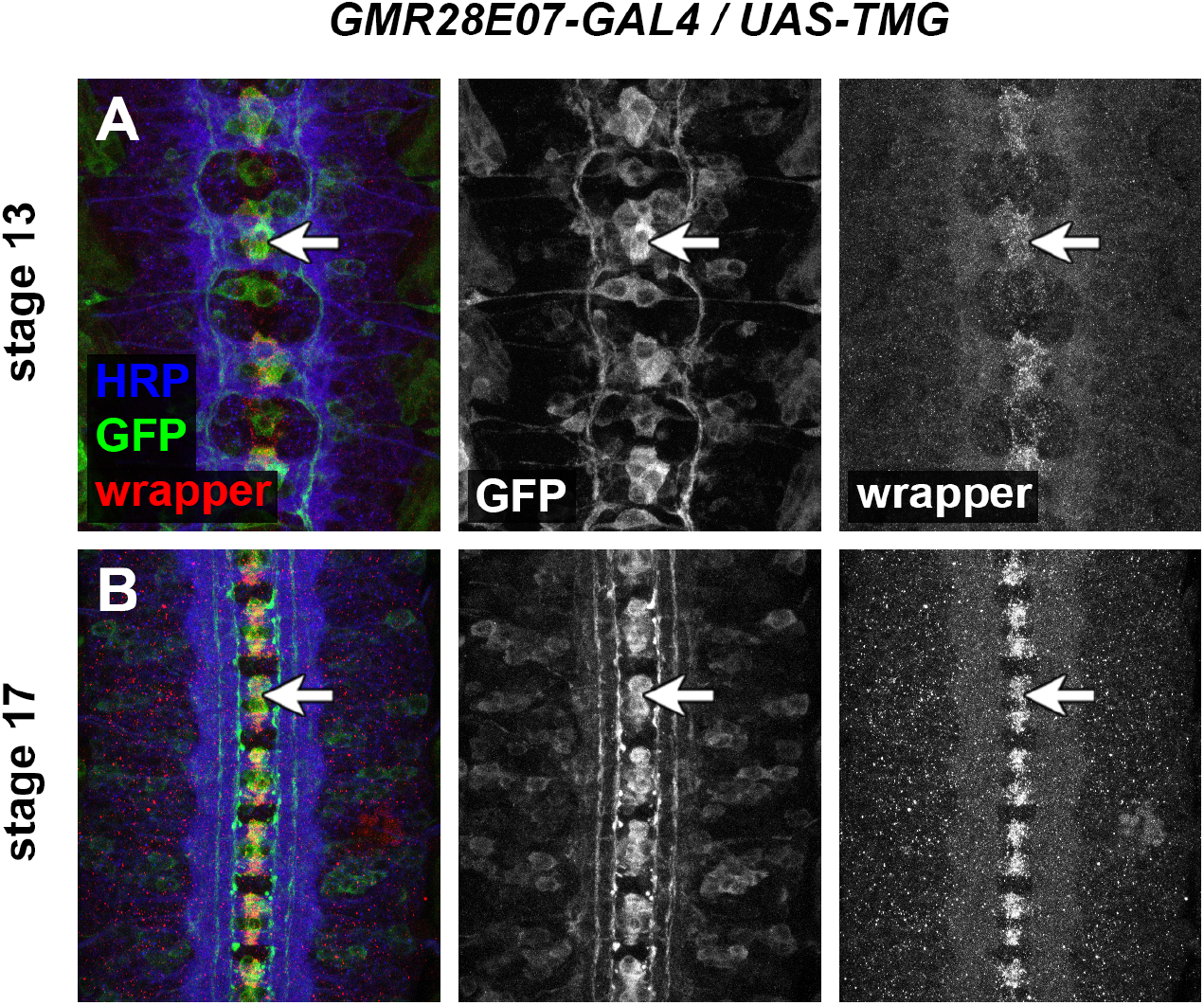
GMR28E07 is expressed in wrapper-positive midline glia. Ventral nerve cords from stage 13 (A) or stage 17 (B) *GMR28E07/UAS-TMG* embryos stained with anti-GFP (green), anti-wrapper (red) and anti-HRP (blue) antibodies. Isolated channels for anti-GFP and anti-wrapper are also shown. Wrapper is a marker for midline glia and is detectable in these cells by stage 12 (Noordermeer *et al.* 1998). At stage 13 (A), GFP expression driven by the *GMR28E07* GAL4 line co-labels wrapper-positive midline glia (A, arrow), along with additional midline-adjacent cells that are not wrapper-positive, some of which are longitudinal pioneer neurons whose axons can be seen extending between adjacent segments. Both GFP and wrapper expression remain strong in the midline glia through stage 17 (B, arrow).

#### Enhancer fragments promoting *robo2* expression in morphologically distinct and genetically separable subsets of lateral longitudinal axons

A number of the putative enhancer fragments presented here exhibit broad expression in the embryonic ventral nerve cord which may overlap with Robo2’s endogenous expression pattern in lateral longitudinal axons. Two fragments in particular exhibit more restricted expression where GFP is specifically detectable in longitudinal axons in the lateral zone: GMR28F02 and GMR28G05 (**Figures 10 and 13)**. These two fragments are non-overlapping **(Figure 1)**, and their GFP expression patterns do not appear to be identical, suggesting that they may represent (at least) two molecularly distinct enhancer regions promoting expression in two distinct subsets of longitudinal neurons. Both subsets include early-crossing commissural axons, with GFP-positive axons visible crossing midline as early as stage 13 in both lines, and both include some GFP-positive longitudinal axons in pathways outside of the lateral zone. As Robo2 protein is not normally detectable on medial/intermediate longitudinal axons at late stages, it is possible that GFP-positive axons visible in the medial/intermediate zones at stage 17 may be due to perdurance of early GFP expression in medial/intermediate pathway pioneer neurons.

To more closely compare the expression patterns of GMR28F02 and GMR28G05, and to determine whether the neurons expressing GAL4 in these two lines represent distinct neuronal subsets, we first compared the cell body positions of the neuron clusters expressing GMR28F02 and GMR28G05 **(Figure 22).** We traced the cell body clusters and the midline-crossing portions of GFP-positive axons in stage 16 *GMR28F02/UAS-TMG* and *GMR28G05/UAS-TMG* embryos **(Figure 22A’, A”, B’, B”)** and overlaid the traces, using the outline of the HRP-positive axon scaffold to register the two traces with each other **(Figure 22C).** We found that the GFP-positive cell body clusters in GMR28F02 and GMR28G05 exhibit little or no overlap in these registered traces. GFP-positive cell bodies in GMR28F02 are found mainly in the anterior portion of each segment, located between the AC of one segment and the PC of the next anterior segment **(Figure 22A, A’, A”**). In contrast, GFP-positive cell bodies in GMR28G05 are clustered near the PC in each segment, with the anterior boundary of each cluster located near the posterior edge of the AC in the same segment **(Figure 22B, B’, B”)**.

**Figure 22.**
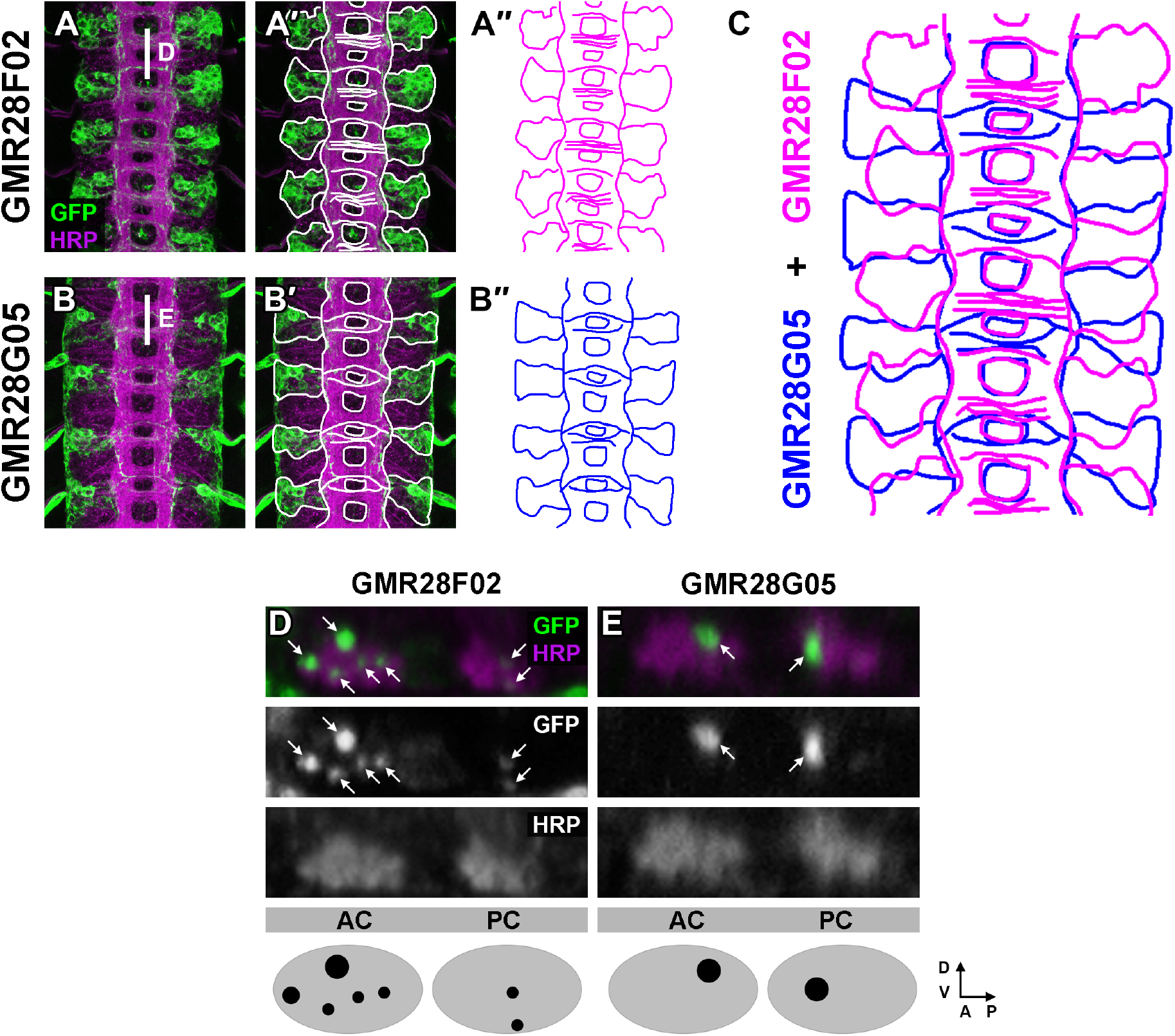
GMR28F02 and GMR28G05 label two distinct subsets of commissural longitudinal neurons. (A,B) Ventral nerve cords from stage 16 *GMR28F02/UAS-TMG* (A) and *GMR28G05/UAS-TMG* (B) embryos stained with anti-GFP (green) and anti-HRP (magenta) antibodies. Anti-GFP labels the cell bodies and axons of neurons that express the indicated GAL4 transgene. White vertical bars indicate the positions of y,z cross-sections shown in (D) and (E). (A’,B’) The same images as in (A) and (B) are shown with the axon scaffold, GFP-positive neuronal cell bodies, and GFP-positive commissural axons traced in white. (A’’,B’’) The tracings shown in (A’) and (B’) are shown in magenta (for GMR28F02) and blue (for GMR28G05). (C) The tracings for the two GAL4 lines are shown overlaid on each other, illustrating that the two GAL4 lines each label distinct clusters of neuronal cell bodies with minimal overlap between their locations in the VNC cortex. The commissural axon tracings suggest that the two subsets of neurons labeled in the two lines also cross the midline at distinct locations within the two commissures. The GMR28G05 tracing has been compressed to 96% of its original height to improve alignment between the two embryos’ axon scaffolds. (D,E) Confocal y,z cross-sections through a single representative segment from the *GMR28F02/UAS-TMG* (D) and *GMR28G05/UAS-TMG* (E) embryos shown in (A) and (B), showing the positions of GFP-positive axons as they cross the midline in the anterior commissure (AC) and posterior commissure (PC). At least five distinct axon bundles are detectable within the AC and two distinct bundles in the PC in *GMR28F02/UAS-TMG* embryos (D, arrows). In *GMR28G05/UAS-TMG* embryos, one main GFP-positive axon bundle each is present in the AC and PC (E, arrows). As sugggested by the axon tracings in (C), the dorsal/ventral and anterior/posterior positions of the commissural bundles suggest that the axons labeled in each line cross the midline at distinct positions from each other. Schematic diagram of the relative sizes and positions of the GFP-positive commissural axon bundles are shown below for the two GAL4 lines. In panels D-E, dorsal is up and anterior is to the left.

We next compared y,z confocal cross-sections through the anterior (AC) and posterior (PC) commissures in these same stage 16 *GMR28F02/UAS-TMG* and *GMR28G05/UAS-TMG* embryos and found that GFP-positive commissural axons in the two subsets cross the midline in distinct locations. GMR28F02 axons cross in one bundle near the posterior edge of the PC and 2-3 bundles in the anterior half of the AC, while GMR28G05 axons cross in one bundle near posterior edge of the AC and 1-2 bundles at anterior edge of the PC **(Figure 22D,E)**, again suggesting that the neurons labeled by these two GAL4 lines represent distinct subsets of cells.

Finally, we used x,z confocal cross-sections to examine the positions of GFP-positive lateral longitudinal axons and FasII-positive axon pathways in stage 16 *GMR28F02/UAS-TMG* and *GMR28G05/UAS-TMG* embryos, compared with equivalent x,z cross-sections in *robo2^myc-robo2^* embryos **(Figure 23).** We confirmed that myc-tagged endogenous Robo2 protein overlaps with lateral FasII-positive longitudinal pathways in stage 16 *robo2^myc-robo2^* embryos, as predicted by previous studies which examined Robo2 and FasII expression in different embryos **(Figure 23A, A’)** (Rajagopalan *et al.* 2000b; Simpson *et al.* 2000b). However, we found that GFP-positive longitudinal axons in *GMR28F02/UAS-TMG* and *GMR28G05/UAS-TMG* embryos did not overlap with lateral FasII pathways **(Figure 23B, B’, C, C’)**, suggesting that these two subsets of Robo2-expressing axons are not among the subset of lateral longitudinal axons that are predicted to be both Robo2-positive and FasII-positive.

**Figure 23.**
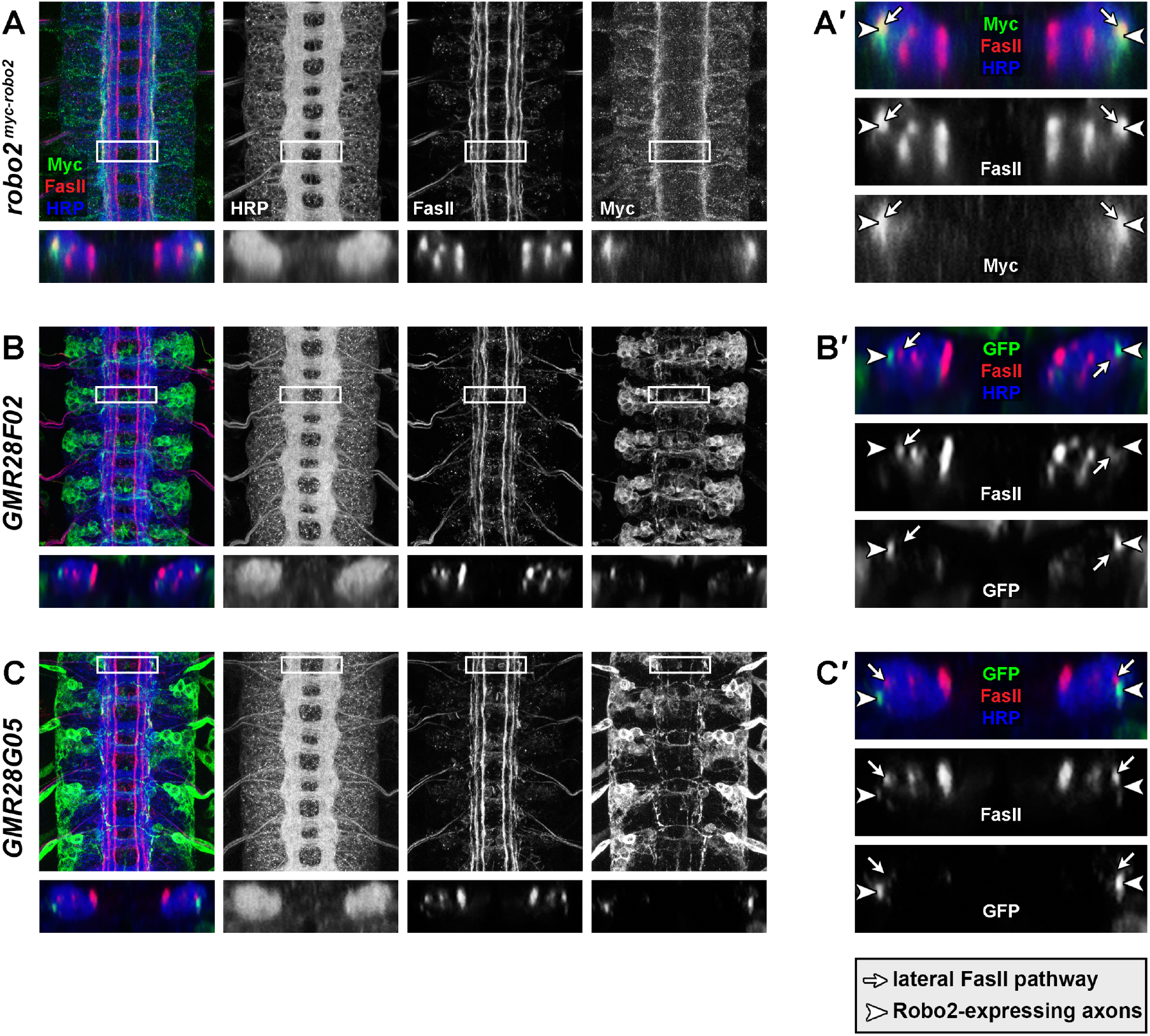
Lateral longitudinal axons labeled by GMR28F02 and GMR28G05 are distinct from FasII-positive lateral axon pathways. (A) Stage 16 *robo2^myc-robo2^* embryo stained with anti-myc (green), anti-FasII (red) and anti-HRP (blue) antibodies. Individual channels are shown in grayscale. Lower panel shows confocal x,z cross-sections through the area outlined by the white box. (B,C) Stage 16 *GMR28F02/UAS-TMG* (B) and *GMR28G05/UAS-TMG* (C) embryos stained with anti-GFP (green), anti-FasII (red) and anti-HRP (blue) antibodies. Lower panels show confocal x,z cross-sections through the area outlined by the white box. (A’-C’) Enlarged view of the confocal x,z cross-sections from panels (A-C). Arrows indicate the position of the dorsal lateral FasII axon pathway, and arrowheads indicate the position of myc-positive (A’) or GFP-positive (B’,C’) axons. While myc-positive axons in A’ overlap with FasII-positive lateral axon pathways, GFP-positive axons in B’ and C’ do not overlap with FasII, indicating that *GMR28F02* and *GMR28G05* label lateral longitudinal axons that are not part of FasII-positive pathways.

## Discussion

Among the three *Drosophila robos, robo2* exhibits the broadest and most dynamic expression pattern and the most diverse set of molecular and developmental roles during embryonic development. Accordingly, the transcriptional regulation of *robo2* expression appears to be more complex than for *robo1*. We and others have constructed *robo1* rescue transgenes containing less than 5 kb of flanking genomic sequence that recapitulate *robo1*’s full expression pattern in the embryonic CNS (Kidd *et al.* 1998b; Spitzweck *et al.* 2010; Brown *et al.* 2015). However, our attempts to create an equivalent rescue transgene for *robo2* with similarly sized flanking sequences have not been successful, suggesting that the regulatory sequences necessary for proper *robo2* expression may be more numerous and/or located farther from the *robo2* transcription start site than for *robo1* (T.A.E., unpublished data). Notably, the *robo2* transcription unit is approximately 40 kb in size, nearly five times the size of *robo1* (~ 8,400 bp) and unlike *robo1* includes a very large first intron (22 kb), which could potentially contain regulatory sequences in addition to any located upstream of the *robo2* promoter.

Here, we identify enhancer regions located upstream of the *robo2* promoter and in the *robo2* first intron which can drive GAL4 expression in specific cell types where *robo2* is known to function, including early pioneer neurons, midline glia, and lateral longitudinal neurons. The identification of fragments containing regulatory sequences described here should enable similar rescue constructs for *robo2,* and may allow rescue of specific phenotypes/subsets of neurons (e.g. restoring *robo2* expression in lateral longitudinal neurons with the GMR28G05 or GMR28F02 fragments, alone or in combination, may allow specific rescue of *robo2’s* role in lateral pathway formation). Our results here also provide insight into the regulatory logic of *robo2* expression, by identifying genetically separable enhancer regions that individually specify expression in distinct subsets of *robo2-*expressing cells (e.g. midline glia vs lateral longitudinal neurons).

### Differences between Robo2 protein expression and GAL4-induced GFP expression

Importantly, the initation and termination of *GAL4* transcription in the transgenic lines described here should closely reproduce the temporal dynamics of transcription specified by each enhancer fragment, but the GAL4 and/or TauMycGFP proteins are likely to perdure well after transcription ceases. Thus, cells that express a given enhancer early during development are likely to remain GFP-positive through later stages, even if the enhancer is no longer active. For this reason, the GFP expression patterns we describe at later stages of development are likely to reflect a summation of each enhancer’s expression throughout all stages of embryogenesis.

The subcellular localization of TauMycGFP will likely also be distinct from Robo2 protein. Cell bodies and the entire length of axons are labeled by TauMycGFP, while Robo2 is primarily localized to axons and is likely not uniformly expressed along the entire length of all axons. For example, the lateral longitudinal neurons labeled by GMR28G05 and GMR28F02 are clearly commissural neurons and the midline-crossing portions of their axons are labeled by TauMycGFP **(Figures 10 and 13)**, but there are few or no commissural axon segments with detectable Robo2 protein at any stage in *robo2^myc-robo2^* or *robo2^HA-robo2^* embryos **(Figure 2)**. The endosomal sorting receptor Commissureless (Comm) prevents Robo1 protein from reaching the growth cone surface as commissural axons grow toward and across the midline, which accounts for the exclusion of Robo1 protein from commissural axon segments (Kidd *et al.* 1998a; Keleman *et al.* 2002, 2005). In *comm* gain of function embryos, Robo2 protein is also downregulated, but it is not clear whether Comm normally regulates Robo2 protein the same way it does Robo1 (Rajagopalan *et al.* 2000a; Simpson *et al.* 2000a).

### Dispensability of *robo2* enhancer elements in introns 2-13

We note that several of the tested fragments located within *robo2* introns 2-13 (GMR27G11, GMR27F07, GMR28D05, GMR27D10, and GMR27D07) were deleted in a series of knock-in and CRISPR-modified alleles created by us and others in which exons 2-14 of *robo2* along with the intervening introns were replaced with cDNAs encoding various *robo* variants, including fulllength *robo2* coding sequences (Spitzweck *et al.* 2010; Howard *et al.* 2021). The *robo2^myc-robo2^* allele reported here also deletes these intronic sequences. We have shown here that GMR27F07, GMR28D05, and GMR27D07 can promote expression in the embryonic ventral nerve cord **(Figure 1).** However, in both our and Spitzweck et al.’s *robo2^robo2^* alleles, introns 2-13 were deleted without any detectable effect on the midline repulsion or lateral positioning activities of *robo2.* Santiago and colleagues also detected no defects in *robo2-*dependent motor neuron innervation of ventral muscles 6/7 in animals homozygous for the *robo2^robo2^* knock-in allele (Santiago *et al.* 2014). These results suggest that any expression conferred by these fragments is dispensable for those axon guidance activities, at least to the extent that can be measured by examining FasII-positive axons in the embryonic ventral nerve cord. It may be the case that the expression patterns conferred by enhancers within introns 2-13 are redundant with patterns conferred by other enhancers that were not deleted in the modified alleles. Alternatively, the ventral nerve cord expression patterns conferred by these enhancers may be necessary for axon guidance or other developmental outcomes that were not examined in the studies cited above. Importantly, we have not examined all *robo2*-dependent developmental contexts in these gene replacement backgrounds, and thus cannot rule out the possibility that introns 2-13 and any associated enhancer elements may be required for some of *robo2’s* developmental roles.

### Developmental timing of Robo2-positive vs FasII-positive longitudinal axon pathway formation

We have shown here that enhancers within the GMR28F02 and GMR28G05 fragments drive expression in distinct subsets of lateral longitudinal neurons. We note that GFP-positive longitudinally-projecting axons are present in the lateral region of the neuropile in *GMR28F02/UAS-TMG* embryos by stage 13, and form a continuous lateral pathway well before the lateral FasII-positive pathways begin forming **(Figure 10C,D).** Myc-tagged Robo2 protein is also detectable within the lateral neuropile in *robo2^myc-robo2^* embryos by stage 13 **(Figure 2C)**, consistent with the idea that Robo2-positive lateral axon tracts form before FasII-positive lateral pathways. The x,z cross sections presented in **Figure 23** reveal that at least some of these Robo2-positive tracts remain distinct from FasII-positive pathways through stage 17. GFP-positive lateral axons in *GMR28G05/UAS-TMG* embryos are also distinct from FasII lateral pathways in cross-sections, but GMR28G05 axons reach lateral positions later than GMR28F02 and are not detectable in the lateral zone until FasII-positive lateral pathways begin forming, around stage 15 **(Figure 13E).**

We and others have previously shown that the lateral positions of Robo2-expressing axons are not affected in *robo2^robo1^* or *robo2^robo2ΔIg1^* embryos, while lateral FasII pathways fail to form correctly in these embryos (Spitzweck *et al.* 2010; Howard *et al.* 2021). Robo2 protein expression appears to overlap with lateral FasII pathways in *robo2^myc-robo2^* embryos, and the generally accepted model of Robo2’s lateral positioning activity is that it guides FasII-positive axons to the lateral region by acting cell-autonomously as a Slit receptor. However, it remains unclear which, if any, FasII-positive lateral axons also express Robo2. Our results suggest that at least some subsets of Robo2-positive lateral axons are distinct from FasII-positive lateral pathways. The GMR28F02 and GMR28G05 enhancer fragments described here provide a method for labeling Robo2-expressing axons independently of their Robo2 expression. This should allow us to directly examine which (if any) of these subsets depend on Robo2 expression for their guidance to the lateral zone, which may help clarify whether Robo2’s function in promoting lateral pathway formation is a cell-autonomous or non-autonomous function.

### Locations of specific enhancer element(s) within each identified enhancer region

Most of the fragments tested here are quite large (3kb) relative to the average size of a single enhancer element or transcription factor binding site. Importantly, our analysis here does not identify specific transcription factor binding sites within any of the identified enhancer regions, or reveal how many sites are present in each fragment, or their specific location(s). In some cases, the overlapping nature of the tested fragments may help narrow down the likely location of specific enhancer regions. For example, fragments GMR28A12 and GMR28F02 overlap with each other and both exhbit similar strong expression in the visceral mesoderm, suggesting that the relevant enhancer sequence(s) for this expression may be located within the 1,020 bp overlap region shared by these two fragments. In contrast, the neuronal expression of GMR28F02 is not shared by GMR28A12, suggesting that any enhancer(s) active in lateral longitudinal neurons are likely to be located within the 2,715 bp region of GMR28F02 that does not overlap with GMR28A12. It is more difficult to determine whether GMR28F02’s lateral axon expression is shared by GMR28E10, which also overlaps with GMR28F02 but has much broader neuronal expression. Similarly, it is difficult to determine if GMR28G05’s lateral neuron expression is shared by its overlapping fragments GMR28D10 or GMR28D12, because the expression patterns of these two fragments are also very broad. Further studies will be necessary to sub-divide each of these regions to determine the number, identity, and location of specific enhancer elements within each.

## Materials and methods

### Drosophila strains

Transgenic *Drosophila* strains carrying GAL4 transgenes from the FlyLight collection (Pfeiffer *et al.* 2008; Jenett *et al.* 2012) were obtained from Bloomington *Drosophila* Stock Center. The following *Drosophila* mutant alleles and transgenes were used: *w^1118^; sna^Sco^/CyO,P{en1}wg^en11^ (Sco/CyOwg), y^1^ M{w[+mC]=nos-Cas9.P}ZH-2A w* (nos-Cas9.P)* (Port *et al.* 2014), *robo2^myc-robo2^* (this study), *robo2^HA-robo2^* (Howard *et al.* 2021), *P{GMR27G11-GAL4}attP2, P{GMR27F07-GAL4}attP2, P{GMR28D05-GAL4}attP2, P{GMR27D10-GAL4}attP2, P{GMR27D07-GAL4}attP2, P{GMR28A12-GAL4}attP2, P{GMR28F02-GAL4}attP2, P{GMR28E10-GAL4}attP2, P{GMR28D10-GAL4}attP2, P{GMR28G05-GAL4}attP2, P{GMR28D12-GAL4}attP2, P{GMR28F12-GAL4}attP2, P{GMR27H02-GAL4}attP2, P{GMR28C04-GAL4}attP2, P{GMR28B05-GAL4}attP2, P{GMR28A10-GAL4}attP2, P{GMR28E07-GAL4}attP2, P{GawB}NP6273 (robo2^GAL4^*) (Hayashi *et al.* 2002), *P{UAS-TauMycGFP}III.* The genomic sequence coordinates for the DNA fragments associated with the above FlyLight GAL4 transgenes are described in **Table 1.**

### Generation of myc-tagged *robo2^myc-robo2^* allele

CRISPR modification of *robo2* was carried out as described in (Howard *et al.* 2021), with an N-terminal 4xMyc tag replacing the 4xHA tag in the full-length *robo2^robo2^* donor plasmid. *robo2* gRNA sequences (GTATCTTTTATACCATGCCA and GGAGCAAAGGTCAGACATTA) were cloned into the tandem expression vector pCFD4 (Port *et al.* 2014) via PCR followed by Gibson assembly using the PCR product and BbsI-digested pCFD4 backbone. The *robo2* gRNA plasmid and homologous donor plasmid were co-injected into *nos-Cas9.P* embryos (Bloomington Drosophila Stock Center stock #54591) (Port *et al.* 2014) by BestGene Inc (Chino Hills, CA). Injected individuals (G0) were crossed as adults to Sco/CyOwg. Founders (G0 flies producing F1 progeny carrying modified robo2 alleles) were identified by testing two pools of three F1 females per G0 cross by genomic PCR. From each identified founder, 5-10 F1 males were then crossed individually to Sco/CyOwg virgin females. After three days, the F1 males were removed from the crosses and tested by PCR to determine if they carried the modified allele. F2 flies from positive F1 crosses were used to generate balanced stocks, and the modified alleles were fully sequenced by amplifying the entire modified locus from genomic DNA, then sequencing the PCR product after cloning via CloneJET PCR cloning kit (Thermo Scientific).

### Embryo collection, antibody staining/immunofluorescence, and imaging

*Drosophila* embryo collection, fixation, and antibody staining were carried out as described by Patel (Patel 1994). Flies carrying each *GAL4* transgene were mated with flies carrying a *UAS-TauMycGFP* transgene, and offspring from the crosses were collected as embryos every 24 hours for 4-5 days. 0-24 hour old embryos were collected on apple juice agar plates containing a dollop of yeast paste (50/50 mix by volume of water and dried active baker’s yeast). A 50% bleach solution was applied directly to the embryos on the agar plate for 2-5 minutes to dissolve the embryonic chorion, then embryos were collected in mesh baskets and rinsed well with water. Dechorionated embryos were transferred to glass scintillation vials containing 5 ml heptane and 5 ml 3.7% formaldehyde and fixed for 12-15 min at room temperature while rocking on a nutator. After fixation, the formaldehyde solution was removed and 10 ml methanol was added, followed by vigorous shaking for 30 seconds to remove embryonic vitelline membranes. Devitellinized embryos were rinsed well with methanol to remove residual heptane and formaldehyde, transferred to 1.5 ml microcentrifuge tubes, and stored in methanol at −20°C. For antibody staining, embryos were rehydrated in PBS + 0.1% Triton X-100 (PBT), blocked with PBT + 5% normal goat serum (NGS), incubated with primary antibodies diluted in PBT+NGS at 4°C overnight, washed in PBT, incubated with secondary antibodies diluted in PBT+NGS for 1-2 hours at room temperature, then washed again in PBT. Embryos stained with HRP-conjugated antibodies were developed by incubation with Stable Diaminobenzidine (DAB) solution (Invitrogen #750118) according to the manufacturer’s instructions. Stained embryos were stored in 70% glycerol/PBS at room temperature. Whole embryo images were acquired using a Leica M165FC stereo microscope with attached MC170 digital camera and processed by Adobe Photoshop software. Ventral nerve cords from fluorescently stained embryos of the desired genotype and developmental stage were dissected and mounted in 70% glycerol/PBS. Fluorescent confocal stacks were collected using a Leica SP5 confocal microscope and processed by Fiji/ImageJ (Schindelin *et al.* 2012) and Adobe Photoshop software.

The following antibodies were used: rabbit anti-GFP (Invitrogen #A11122, 1:1000), rabbit anti-c-Myc (Sigma #C3956), mouse anti-Fasciclin II (Developmental Studies Hybridoma Bank [DSHB] #1D4, 1:100), mouse anti-HA (BioLegend #901503, 1:1000), mouse anti-wrapper (DSHB #10D3, 1:100), HRP-conjugated goat anti-mouse (Jackson ImmunoResearch #115-035-003, 1:500), HRP-conjugated goat anti-rabbit (Jackson #111-035-003, 1:500), Alexa 488-conjugated goat anti-HRP (Jackson #123-545-021, 1:200), Alexa 647-conjugated goat anti-HRP (Jackson #123-605-021, 1:100), Cy3-conjugated goat anti-mouse (Jackson #115-165-003, 1:1000), Cy3-conjugated goat anti-rabbit (Jackson #111-165-003, 1:500), Alexa 647-conjugated goat anti-mouse (Jackson #115-605-003, 1:500), Alexa 488-conjugated goat anti-rabbit (Jackson #111-545-003, 1:500), Alexa 647-conjugated goat anti-rabbit (Jackson #111-605-144, 1:500).

## Acknowledgments

Stocks obtained from the Bloomington Drosophila Stock Center (National Institutes of Health [NIH] grant P40 OD-018537) were used in this study. Monoclonal antibodies were obtained from the Developmental Studies Hybridoma Bank, created by the Eunice Kennedy Shriver National Institute of Child Health and Human Development of the NIH and maintained at The Department of Biology, University of Iowa, Iowa City, IA 52242. This work was supported by NIH grant R15 NS-098406 (T.A.E.).

## Notes

### Competing Interest Statement

The authors have declared no competing interest.

